# Assessment of variant effect predictors unveils variants difficulty as a critical performance indicator

**DOI:** 10.1101/2024.07.08.602580

**Authors:** Ragousandirane Radjasandirane, Julien Diharce, Jean-Christophe Gelly, Alexandre G. de Brevern

## Abstract

Amino acid substitutions in protein sequences are generally harmless, but a certain number of these changes can lead to disease. Accurate prediction of the impact of genetic variants is crucial for clinicians as it accelerates the diagnosis of patients with missense variants associated with health issues. Numerous computational tools have been developed for prediction of the pathogenicity of genetic variants based on different methodologies. Nowadays, many approaches are based on Machine Learning. Assessment of the performance of these diverse computational tools is crucial to provide guidance to both future users and especially clinicians. In this study, a large-scale study of 65 tools was conducted. Variants from both clinical and functional context have been used, incorporating data from the ClinVar database and bibliographic sources. The analysis showed that AlphaMissense is often performing very well and is actually the best option among existing tools. Additionally, meta-predictors, as expected, are of high quality and perform well on average. Tools using evolution information demonstrated highest performances on functional variants. These results also highlighted some variations in the difficulty to predict some specific variants while others are always well categorized. Strikingly, the majority of variants from the ClinVar database appear to be easy to predict, while variants from other sources of data are more challenging. These results demonstrate that this variant predictability can be classified into three distinct classes: easy, moderate and hard to predict. We analyzed the parameters leading to these differences and show that classes are linked to structural and functional information.

## Introduction

In living organisms, DNA molecules are subject to replication cycles, a process fundamental for the generation of new cells and the transfer of genetic information. Sometimes, alterations occur in the nucleotide sequence during duplications, leading to different outcomes. In the first case, mutations are neutral, having no effect on the cell and the biological processes. The second one occurs when nucleotide change is important enough to impact cells adversely, leading to disease. Finally, the most severe case involves a nucleotide alteration so fatal that the cell cannot survive^1^. These nucleotide changes can range from a single nucleotide variation (Single Nucleotide Polymorphism or SNP), to larger deletions / insertions, frameshifts or even break and rearrangements of a large portion of sequence. In this study, we focus on SNPs, *i*.*e*. punctual change of nucleotides in humans. The sequencing of several human genomes has revealed the extensive nature of human polymorphism, indicating that most SNPs are harmless. Only a fraction of them have been directly linked to diseases^2^. The first set of SNPs are called benign, the second pathogenic.

The criterias for deciding whether a variant should be annotated as benign or pathogenic are now well established. A recognized guideline is provided by the American College of Medical Genetics (ACMG) and Association for Molecular Pathology (AMP)^3^, it lists several good practices to classify newly sequenced variants. These variants are cataloged in specific databases such as dbSNP^4^, OMIM^5^ or Human Gene Mutation Database (HGMD)^6^. The best known being ClinVar^7^, a clinical database associating the variant sequences with the related disease (if available) and their clinical significance (*e.g*., benign, pathogenic or uncertain significance). This database links the relationship between phenotype and human polymorphisms. As variants can be found when sequencing patient samples, it is also important to provide *in vitro* tests to assess their impacts; these experimental tests are often associated to *in silico* evaluations, *e.g*.,^8,9^. However, conducting experiments to systematically assess the impact of variants is often both time consuming and expensive.

Hence, since the 2000’s, software-based prediction of variant effects have emerged, enabling fast and free evaluation of variant impact without going through an experimental validation phase^10^. These methods are called Variant Effect Predictors (VEPs). The characterization of the variant effects could be very useful in diagnosing less known diseases, in particular for clinicians. The impact of such approaches has been phenomenal. Two pioneering approaches stand out: Sorting Intolerant from Tolerant (SIFT)^11^ and PolyPhen^12^, accumulating over 15,000 citations each by 2024, demonstrating their appeal.

VEPs can be divided into several types depending on their methodologies. The first group of VEPs were based on the simple principle of the conservation of amino acids through evolution, with the idea that mutations in conserved positions have more chance to be deleterious than one on less conserved sites^13^. The impact can be greater if the physco-chemical properties of the initial residue are not conserved as observed for mutation matrices^14^. The most known VEPs using this approach are SIFT and PolyPhen2, but many others have been developed such as PROVEAN^15^ or DEOGEN2^16^. The second group of VEPs have integrated machine learning (ML) to evolutionary information previously described (*e.g.*, SuSPeCT^17^, PrimateAI^18^, EVE^19^ or CPT^20^). This group also comprises AlphaMissense^21^ developed by DeepMind (from Alphabet) who adapted AlphaFold2 model^22^ to the problematic of predicting impact of missense variants. Recent evaluations of AlphaMissense show an ambiguous conclusion about its performance where some show a good performance but with modest improvement^23,24^, while other research have demonstrated its high accuracy on real-life data^25^. The third group called “meta-methods” are algorithms using different predictions from published tools as input features; they are generally more accurate than methods incorporated in their algorithms taken alone as demonstrated by tools like BayesDel^26^ or Eigen^27^. Recently, with the rise of Artificial Intelligence (AI) and more specifically deep learning (DL) approaches, the fourth group has been developed by combining meta-methods and ML or AI, which may increase once more the performance, like MetaRNN^28^ which uses a neural network combined with functional scores from 16 existing softwares and 8 conservation scores to produce the prediction.

The majority of the above VEPs are supervised, meaning that their algorithms need to be trained on data with annotations (*e.g.,* benign or pathogenic) in order to accurately predict back these annotations by learning variant characteristics associated with them. Supervised VEPs are sensitive to data issues such as incorrect annotations which lead the VEP to replicate these incorrect annotations, redundancy or other bias potentially present in the dataset used for training^29^. Other VEPs are based on unsupervised methods. These algorithms do not depend on traditional variant annotations. Instead, they use information that is indirectly related to these annotations for making predictions. These VEPs may not have a specific training process and can instead be based on algorithms that capture specific information useful for prediction of variant annotations. The best-known example in this category of non-trained unsupervised VEP is SIFT as it uses evolutionary information through residue conservation although this information is not directly related to variant annotations to predict. EVE is another example of unsupervised VEP trained on capturing evolutionary information through Multiple Sequence Alignment (MSA) using deep learning^19^.

Around a hundred VEP tools have been published^10,30^ with a dramatic increase this past decade. This large number makes it difficult for non-specialist users, such as clinicians (and even specialists), to choose which VEP to use. Selecting reliable tools to quickly assess the impact of a new variant is clearly a real challenge. It is crucial to have an independent benchmark of these VEPs to highlight the most relevant ones based on their performance.

One of the major drawbacks of benchmarking a high number of methods is to prepare unbiased datasets. It is essential not to have any variants that have been taken into account in the development of the methods tested. Several common biases exist, they are referred to as data circularity^31^. The most common data circularity occurs when the evaluation includes data that has already been used to build the model being tested. Other types of data circularity will also be discussed in this study.

Recently published benchmarks are managing these biases either by using independent variant annotations unused by VEPs^31,32^ or data from Deep Mutation Scanning (DMS) experiment^30^. The main limitation of such benchmarks is the number of VEPs tested which is often restricted, except in Livesey et al.^30^, where authors assessed 55 VEPs. Additionally, the reasons behind the failures of VEPs remain unclear and insufficiently discussed in those papers.

Here, a new exhaustive benchmark of a large set of 65 VEPs across three different datasets was performed, with variants that are clinically relevant or are functionally impacting. Each dataset has been rigorously processed to reduce, as much as possible, biases that could inflate performances of evaluated VEPs. Our results show that VEPs perform differently depending on whether they are tested on clinical variants or functional ones. We also emphasize the simplicity of use, because a powerful but complex tool would not be useful to healthcare professionals, who are not specialists in bioinformatics tools and their manipulations. An in-depth analysis of the results obtained for each dataset reveals that an unexpected parameter has a crucial importance: the prediction difficulty. It means that certain variants have their level of pathogenicity easily predictable by almost all tools, while for some others, the majority of VEPs yield incorrect predictions. Consequently, the variants were clustered in three new classes: easy, moderate, and hard to predict. We demonstrate that this classification is relevant regarding characteristics related to the structure of the protein and its function.

## Materials and Methods

### 1.1. Datasets

For our present study, clinical variants were retrieved from two main sources: the ClinVar database^7^ and a “Clinical dataset” published in^33^ were used. Additionally, we used a functionally characterized dataset based on UniProt mutagenesis data named “UniFun” which is adapted from^32^

#### 1.1.1. ClinVar

The first dataset is taken from the ClinVar database, the reference database listing the genetic variants and their pathogenicity. Only variants recorded in the database after 2021, May 1^st^ were kept as numerous methods have been trained on ClinVar prior to May 2021. Selected variants were obtained with filters such as: “Benign”, “Likely benign”, “Likely pathogenic”, “Pathogenic”, “Missense” and “Single nucleotide” terms in ClinVar. Labels “Likely benign” are considered as “Benign” and labels “Likely pathogenic” are considered as “Pathogenic” as commonly done^34,35^.

ClinVar separates variants into 4 groups: variants that have (i) at least one submission, (ii) have exactly one submission, (iii) have multiple submissions, and (iv) have been reviewed by an expert panel, suggesting that the annotation is of high quality. This gradient of quality might have an impact on the observed performances. We therefore separated ClinVar performances into two groups: (i) whole ClinVar and (ii) high quality variants that regroups variants have been submitted multiple times or reviewed by an expert panel. This last dataset is called ClinVar_HQ.

#### 1.1.2. Clinical

The second dataset is the Clinical dataset, retrieved from^33^. It does not come directly from a public database, but emerges from a combination of three different sources: (i) the Deciphering Developmental Disorders (DDD) study^36^, a UK-wide collaborative project that aims to unveil the genetic architecture of developmental disorders, (ii) Sanger sequencing and Next Generation Sequencing from clinical diagnostic laboratory^37^, and (iii) Exome sequencing of Amish individuals from a Community Genomics research study^33^. This dataset contains high quality variants that have passed numerous variant filters from clinicians routines and were manually annotated.

#### 1.1.3. UniFun

Our last dataset differs significantly from the others as it is composed of functional variants from mutagenesis experiments reported in UniProt^38^, and not clinical variants. This dataset was initially built from the mining of Uniprot annotations associated with experimentally tested variants^39^. The missense variants were annotated as “benign-like” if the description of the associated mutagenesis experiment had keywords such as *“No effect on function/activity”,* and annotated as “pathogenic-like” if the keywords found are *“Loss of activity/Abolish function/Loss of function/Abolish of interaction”.* However, description of some variants could sometimes be ambiguous, leading to a double “benign-like” and “pathogenic-like” annotation. For example, the variant Y888V of the gene AP2B1 had the description *“Strongly reduces interaction with SNAP91, EPN1 and clathrin. No effect on EPS15 binding. Abolishes interaction with ARRB1 and with DENND1B”.* By consequence, this variant is annotated both as “benign-like” and “pathogenic-like” according to authors^32^, though the interacting proteins are not the same. Thus, they can introduce biases if they are retained in our dataset. Therefore, we analyzed once again the UniProt data bank using different filters to rebuild this particular dataset with our own annotation criterions. If the description of the mutagenesis experiment contains at least one keyword associated to a potential “pathogenic-like” variant (such as *“reduces”, “strongly”, “impaired”, “decrease*”, *“decreases*”, *“decreased*”, *“abolishes*”, *“reduced*”, *“loss*”, *“impairs*”, “*abolished*”, *“disrupts*”, *“disrupted*”, *“misfolding*”, *“defective*”, *“disruption”*, this variant is labeled as “pathogenic-like”. On the contrary, if the description has only *“No effect on / Does not affect”* labels, the variant is annotated as “benign-like”.

For the purpose of the study and to ensure uniform analysis protocol across all three datasets, we have considered these “benign-like” and “pathogenic-like” as “benign” and “pathogenic” variants for this dataset, while they have not been initially clinically categorized. This will allow us to increase the amount of data available for the evaluation of VEPs, enabling a more comprehensive analysis of their accuracy in processing variants from different sources.

#### 1.1.4. Data preprocessing protocol

Two common biases exist when doing evaluation of models. They are known as data circularity of type 1 and type 2^31^. In our case, type 1 circularity (T1C) concerns the use of variants for evaluation that have already been used to train the evaluated model, which will ultimately inflate the measured performance of the models. T1C in the ClinVar database was taken into account by removing data prior to the date 2021/05/01 since models trained on ClinVar were trained on those anterior data. These eliminated variants were also removed from Clinical and UniFun datasets.

A second filter must also be applied to avoid type 1 circularity due to the Humsavar dataset from UniProt ^38^. Humsavar corresponds to all missense variants reported in UniProt and are annotated as (likely) pathogenic, (likely) benign or uncertain significance. These data may have been used to train several VEPs such as VARITY^40^, which implies that these variants are already known by these VEPs. Even though the major part of VEPs are not using specifically the dataset Humsavar, variants from this dataset might be present in other databases since UniProt is generally a well-known data bank. By consequence, those could have been used to train VEPs. Therefore, variants from Humsavar are removed from all our datasets by principle.

Even after excluding variants potentially used in model training from our evaluation datasets, a more subtle form of bias may persist. If a gene is associated with only one high-frequency class (either benign or pathogenic), the model may tend to predict only this class for that gene, regardless of the intrinsic nature of these new variants. Although these predictions may appear to be accurate and contribute to the overall performance of a VEP, this new prediction is not the result of balanced learning, but simply a bias toward one class for that gene. This phenomenon is known as type 2 circularity bias (T2C). Hence, only genes that have a proportion between 40% and 60% of either benign or pathogenic annotation were kept in our study. Similar solution has previously been used to analyze the effect of T2C on VEP’s performances and it has been shown that T2C falsely improves VEP performance scores if not removed^31^.

Nonetheless, it must be noted that data circularity biases can not be totally removed, particularly given the high number of different tools evaluated here. Each tool has been prepared by different research teams with different specific training datasets (for those that require training process) and details or datasets are not always available.

### 1.2. Softwares

54 different methods have been successfully selected^11,12,15–21,26–28,40–80^. They correspond to 65 prediction scores, as some methods provide multiple predictions, *e*.*g*., VARITY that yields 4 different prediction scores corresponding to 4 different versions of VARITY^40^. These additional scores are also considered as an independent VEP. Each VEP is described in Table 1 which summarizes all the information on these VEPs.

**Table 1.**
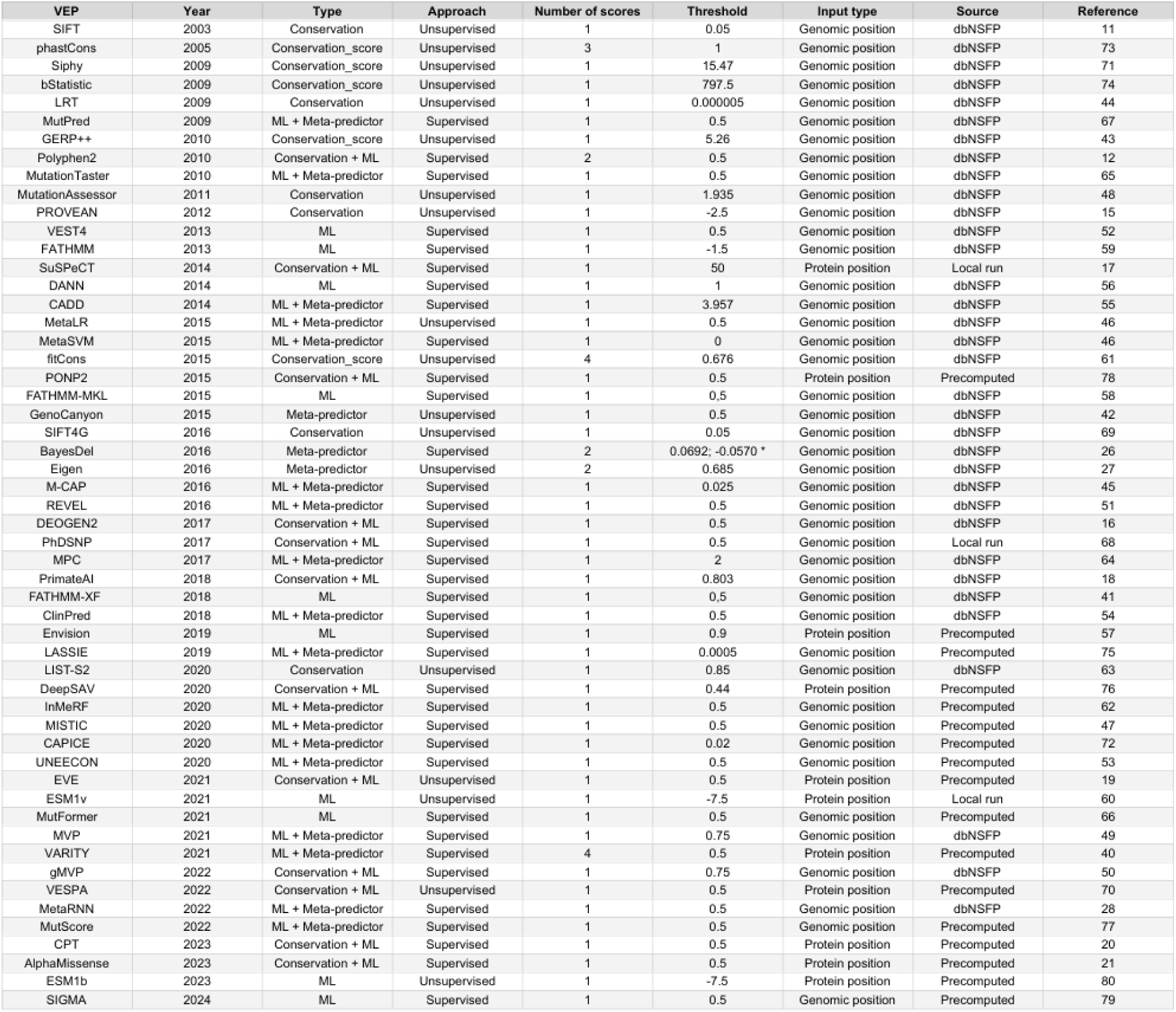
List of the VEPs evaluated in this study. For each VEP is provided its year of publication, the type of the algorithm (6 classes are defined: Conservation, ML, Conservation + ML, Meta-predictor, ML + Meta-predictor, Conservation score, see section Groups of VEP) and approach (supervised or not), the number of scores provided, the threshold used and the source where predictions were retrieved. These last can come from three possible sources. (i) dbNSFP, predictions are coming from the database of prediction dbNSFP, (ii) Precomputed predictions, prediction are coming from parsing files of precomputed predictions provided with the VEP, and (iii) Local run, prediction are coming from local run of the VEP. * BayesDel has two versions and a different threshold for each. BayesDel_addAF 0.0692 and BayseDel_noAF -0.057

Scores of a total of 42 VEPs were retrieved from dbNSFP database version 4.4a^81,82^. Some VEPs have released pre-computed predictions for almost all possible single amino acid substitutions from UniProt human genes, facilitating the score retrieval and thus the analysis. For each of these VEPs, pre-computed prediction files have been downloaded and parsed using homemade python scripts.

Lastly, several VEPs were run locally like PhD-SNPg^68^ and SuSPect^17^ which were run respectively by using python2 and perl scripts provided by authors.

ESM1v^60^ was also run locally using scripts available in the ESM github. For each variant, these scripts generate 5 scores using 5 models and the final score of ESM1v is the mean of generated scores as suggested by authors.

We had to generate several file formats for variant inputs since all VEPs do not require the same data format as input. Some VEPs required input files with genomic positions of variants while other VEPs required protein position instead. TransVar^83^ has been used to switch from one type of position to another and to collect genomic positions from both GR38 and GR37 genome references.

### 1.3. Group of VEPs

We grouped VEP into 6 categories (see Table 1 and Figure S1B) according to their algorithms and features they use. The first group is the “Conservation” group with VEP using evolutionary derived features such as Multiple Sequence Alignment (MSA) to predict the impact of variants based on the conservation of the modified residue. VEP such as SIFT^11^, MutationAssessor^48^ or PROVEAN^15^ are in the “Conservation” group. The second group of VEP uses features similar to those employed by the Conservation group, but integrates Machine Learning algorithms in their model. This group is called “ML + Conservation” and it contains VEP like PolyPhen2^12^, VESPA^70^, EVE^19^ and AlphaMissense^21^.

The third group of VEP, called “ML”, are based on machine learning algorithms but integrate more general features such as allele frequency or the variant amino acid type. This group includes VEST4^52^, MutFormer^66^, ENVISION^57^ and Protein Language Models such as ESM1v^60^ or ESM1b^80^.

The fourth group of VEP combines scores from existing VEP as features to create a new score to predict variant effect. VEP such as Eigen^27^ or BayesDel^26^ are in this group called “Meta-predictor”. These meta-methods can be combined with machine learning algorithms, which constitute our fifth group of VEP called “ML + Meta-predictor”.

Finally, a last group has been created for conservation scores which are scores that can detect evolutionary constrained regions. These scores were not initially developed to detect pathogenic or benign variants but this evolutionary information is commonly used for this purpose especially in meta-methods^26,28,54^, which is why we integrated these scores in the analysis. This group is composed of methods like phastCons^73^, fitCons^61^ or GERP++^43^.

### 1.4. Common metrics

The Area Under the ROC curve (AUROC) metric is commonly used in method evaluation. A ROC curve shows the trade-off between true positive rate (TPR) and false positive rate (FPR) across different decision thresholds. AUROC was obtained here using the “roc_auc_score” function from sklearn python library and by considering pathogenic annotations as positive labels. Some VEPs have an inverted score distribution with lower scores corresponding to benign prediction whereas for the majority of VEP, the lower scores correspond to pathogenic prediction. This inverted distribution affects the AUROC calculation and therefore we simply compute the inverse of the AUROC value for these VEP to get the final AUROC value.

A confusion matrix is generated between true and predicted annotations for each VEP that count the amount of variant correctly predicted in each class (benign or pathogenic). From this matrix, the amount of: (i) pathogenic variants predicted as pathogenic (True Positive or TP), (ii) benign variants predicted as benign (True Negative or TN), (iii) pathogenic variants predicted as benign (False Negative or FN) and (iv) benign variants predicted as pathogenic (False Positive or FP) can be extracted. These are used to compute additional metrics namely sensitivity, specificity, Balanced Error Rate (BER) and the Matthews correlation coefficient (MCC) with following formulas:

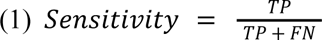

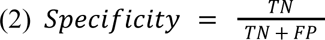

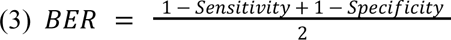

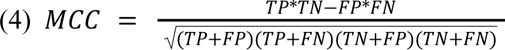

### 1.5. Rank score

Each VEP may have some missing predictions on datasets used here. This raises the issue of VEPs comparison using unshared variants which may produce unrepresentative evaluation. We therefore used a rank score to evaluate each VEPs using shared variants enabling a more comparable evaluation of VEP performances. We have adapted the method from^30^ by adding a statistical test in the comparison.

The rank score is computed by comparing each VEP pair using only shared variants by both VEPs. If predictions of both VEPs are statistically different, one point is given to the winning VEP having the best MCC on shared variants. If predictions are not statistically different, this means that both VEPs are producing similar predictions and it is considered as a tie, each VEP earns 0.5 point. At the end, each VEP has a total of winning points that represent its ability to perform better than other VEP. These points are divided by the total number of comparisons done to produce the final rank score. McNemar statistical test^84^ is used to assess the difference between both predictions through the confusion matrix. This test is available on the python library “statsmodels” with the function “contingency_tables” (see https://www.statsmodels.org/stable/contingency_tables.html).

### 1.6. Error rate per variant

This metric is used to estimate the difficulty of a considered variant to be correctly predicted by our pool of VEPs tools. The error rate per variant metric is computed for each variant by dividing the number of incorrect predictions by the total number of predictions available for this variant, since some VEP may not produce predictions for some variants. This error rate goes from 0 to 1, indicating the relative prediction difficulty for each variant.

### 1.7. Solvent accessibility analysis

Solvent accessibility values were computed using the DSSP program^85^. We analyzed the solvent accessible surface area (SASA) of variants by retrieving AlphaFold2 models from AlphaFoldDB^86^ and experimental structures from Protein Data Bank (PDB)^87^ when available Variant positions having a pLDDT less than 80 were removed from the analysis as the modeled region is not of sufficient quality to interpret the SASA of the residue.

### 1.8. Protein function datasets

To perform an analysis related to protein function, a specific study was carried out using the UniProt data bank. Each gene has been downloaded in Text format, and lines concerning the function have been extracted (line starting with “KW” in the text file). As proteins can be associated with multiple functions, only proteins associated with one major function have been selected. Seven key protein function terms have been selected: (i) Apoptosis, (ii) Immunity, (iii) Metal-binding, (iv) Transport, (v) Signaling, (vi) Enzymatic activity, and (vii) Receptor activity. These groups were selected as they were composed of a sufficient number of variants while having the least overlap with others function terms. All three datasets have been concatenated for each function annotation in order to perform relevant statistical test. Pairwise chi-square test was used to infer the distribution differences between groups while applying a Bonferroni correction as we are testing the same function multiple times. The chi-square test is available in the scipy python library with the module “chi2_contingency” (see https://docs.scipy.org/doc/scipy/reference/generated/scipy.stats.chi2_contingency.html). The Bonferroni correction consists of dividing the alpha threshold by the number of tests done for each group, i.e. 6, giving a corrected alpha of 0.0083 (or 8.3*10^-3).

## Results

### 2.1. Data processing

To avoid data circularity bias of type I (T1C) and type II (T2C) circularity (see section 1.1.4.), successive filters were applied to our datasets (see Figures 1 and S1A).

**Figure 1.**
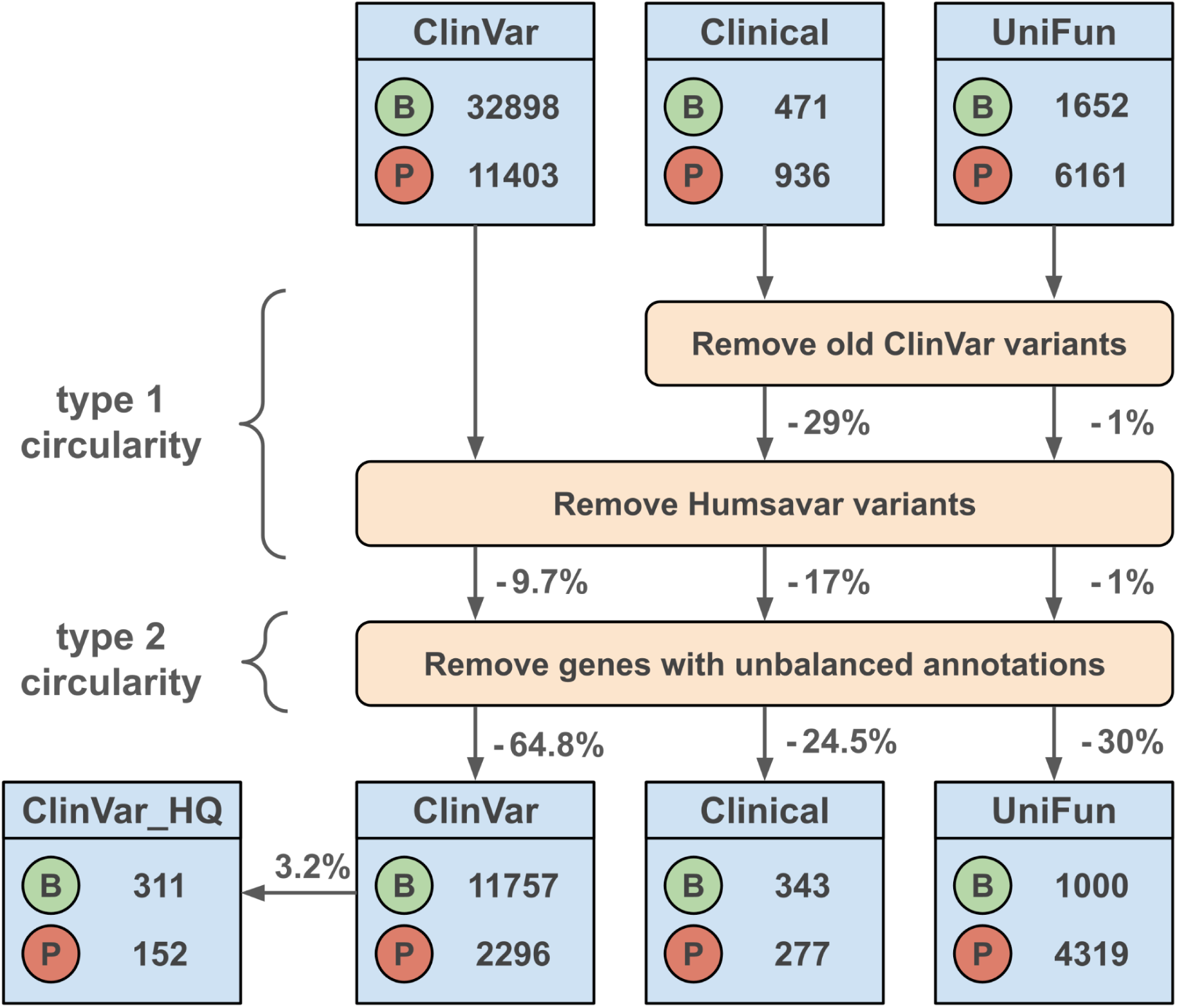
Data processing steps. Each dataset is processed to reduce type 1 and type 2 circularity biases. Benign variants are represented with a green “B”, and pathogenic variants with a red “P”. After each step, the percentage (with minus) of removed data is represented based on the previous pool of data. The percentage (without minus sign) between ClinVar and ClinVar_HQ does not represent a removal but a selection of data from ClinVar. The B and P labels have been kept for UniFun for the sake of simplicity, even though they do not correspond to a real benign and pathogenic state.

The initial ClinVar^7^ dataset corresponds to variants published after 2021/05/01 in ClinVar and encompass 32,898 benign and 11,403 pathogenic variants, *i*.*e*. a total of 44,301 variants. Using these recent variants reduces T1C bias. Similarly, variants present in the Humsavar dataset were removed as this dataset has been used to train numerous VEPs, *e.g.*, SuSPect. 4,347 variants from the initial ClinVar dataset, corresponding to 9.7% of the dataset, were then removed. As described in section 1.1.4., removing genes with unbalanced annotation proportion can greatly help to reduce T2C within the processed dataset. This filter eliminated a significant 65% of the remaining data set, *i*.*e*. 26,020 variants. The final ClinVar dataset thus comprised 14,053 variants, of which 11,757 are benign and 2,296 pathogenic.

From this dataset, a high quality dataset named ClinVar_HQ (High Quality) composed of variants with multiple submissions or reviewed by an expert panel was built. This dataset is made of 311 benign and 150 pathogenic variants and corresponds to 3.2% of the processed ClinVar dataset.

The Clinical dataset has initially 1,407 variants with 471 benign and 936 pathogenic variants. The removal of old variants from ClinVar resulted in the deletion of 419 variants, corresponding to 29% of the initial data set. The cleaning of Humsavar variants removed 169 variants, *i*.*e*. 17% of the dataset. Finally, genes with unbalanced annotation proportion were removed to reduce T2C bias, this last filter has removed 202 variants corresponding to 24.5% of the remaining dataset. The final Clinical dataset is composed of 620 variants with 343 benign and 277 pathogenic variants (see Figure 1).

The UniFun dataset is initially composed of 7,813 variants with 1,652 benign-like and 6,161 pathogenic-like variants. Potential T1C variants correspond to only 105 variants, *i*.*e*. less than 1% of the initial dataset. Similarly, the Humsavar filter has removed only 46 variants of the dataset. The filter to reduce T2C bias eliminated a larger proportion of the dataset by removing 2,381 variants, corresponding to 30% of the dataset. As a result, the cleaned UniFun dataset contains 5,319 variants of which 1000 are benign and 4,319 are pathogenic.

Overall, all these filters have removed 68%, 56% and 30% of respectively the initial pool of ClinVar, Clinical and UniFun dataset in order to reduce type 1 and 2 circularities biases (see Figure 1). However, it must be noticed that data circularity biases can not be totally removed since a high number of VEP are tested, and each of them have specific training datasets (for those that require training process). Furthermore, some variants of these training datasets may still overlap with variants in the refined datasets. It is clearly difficult to create totally unbiased datasets with so many VEPs.

### 2.2 Available predictions

Because of VEP complexity and the large number of data considered, it is not possible to have 100% of the predictions for all the VEPs (which also explains the absence of some tools that we had tried to integrate). Fortunately, many VEP predictions are available for most datasets. Hence, 57 VEPs have a mean data coverage higher than 90% (see Figures S2A and S2B), whereas two VEPs have a mean data coverage of less than 70%, *i*.*e*. SIGMA^79^ and EVE^19^. EVE had the lowest coverage especially for ClinVar and UniFun dataset (see Figure S2B). Its precomputed predictions are mostly incomplete for the Human proteome. We decided to not complete the predictions for EVE as it needs Multiple Sequence Alignment (MSA) and training the model for each of our 9,000 genes is highly time-consuming. The low coverage of EVE is accentuated by our filtering steps that remove a large part of the initial variant (see Figure 1). For this reason, performances of EVE should be carefully considered.

### 2.3. Performances measured by AUROC, MCC and rank score metrics

#### 2.3.1. ClinVar

##### 2.3.1.1. Initial ClinVar dataset

The evaluation of VEPs on the ClinVar dataset gives results of remarkably high quality; almost all VEPs have an AUROC performance value above 0.8 (see Figures 2A and S3A, and Table 2), meaning that they are performing very well at predicting pathogenic class. The top three best VEPs on ClinVar dataset are MetaRNN (AUROC of 0.983), ClinPred (AUROC of 0.980), and BayesDel_addAF (AUROC of 0.977); they are all meta-methods. The best VEPs excluding meta-methods are CPT, VEST4 and AlphaMissense with an AUROC value of 0.965, 0.963 and 0.960 respectively. VEPs that predict slightly better than these latter were initially trained on ClinVar (MetaRNN, ClinPred, BayesDel_addAF and VARITY). Even if the variants potentially used during their training have normally been removed, certain biases could still remain during their training. Indeed, these VEPs may have learned some underlying pattern from the ClinVar database, such as overrepresentation of gene or substitution, that might be also observed in our ClinVar dataset. Despite these potential biases, AlphaMissense, CPT and VEST4 performed similarly as meta-methods with very close AUROC values.

**Figure 2.**
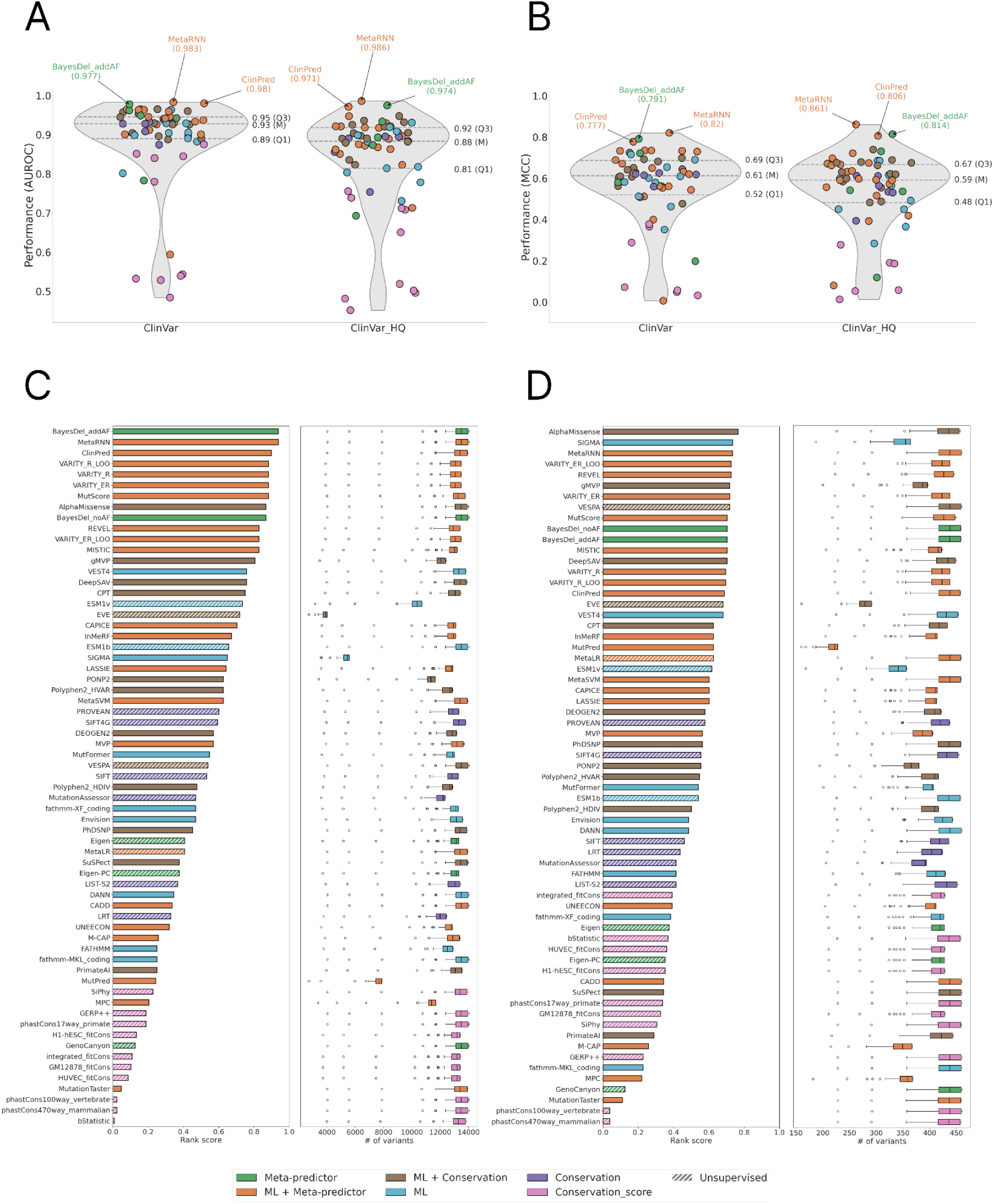
Performances of ClinVar and ClinVar_HQ datasets. (above) Distribution of AUROC (A) and MCC (B) values for ClinVar and ClinVar_HQ datasets. Each dot represents one VEP colored according to the type of algorithm used by the VEP. At the right of each violin plot, first quartile (Q1), median (M) and third quartile values (Q3) of each distribution are represented. The name of the top three VEPs are shown for each dataset. (below) Histogram of rank score values for ClinVar (C) and ClinVar_HQ (D) datasets. Each panel contains rank score values per VEPs on the left, and the distribution of the amount of shared variants used in pairwise comparisons to compute the rank score on the right.

**Table 2.**
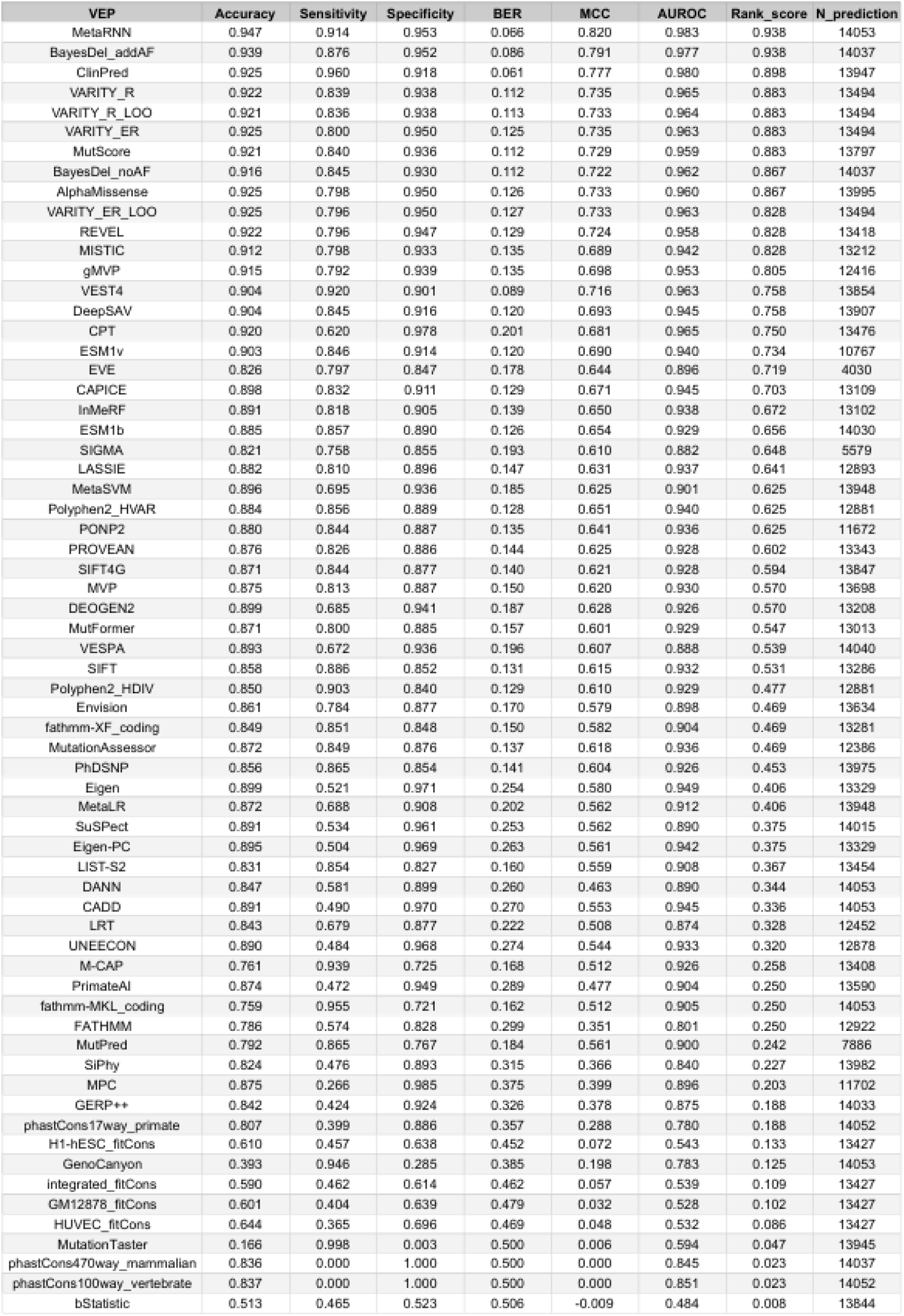
ClinVar performances. Overview of performances of each VEP on ClinVar dataset. Rank score and the number of predictions available are also provided.

The AUROC metric is useful for assessing a model’s ability to correctly classify positive labels (in our case pathogenic class), but does not reflect performance on negative labels (benign class). The MCC metric can undoubtedly better assess the overall performance of tested VEPs. It is clearly seen as MCC value distributions are lower than AUROC values. MetaRNN still performs the best, with a MCC of 0.820 (see Figures 2B and S4A, and Table 2). ClinPred was the second best VEP based on the AUROC values, but is overtaken by BayesDel_addAF with an MCC of 0.791 against 0.777. All four versions of VARITY and AlphaMissense have an MCC of 0.733, these values are 10% lower than the highest MCCs, underlying a larger gap in terms of performance than seen with AUROC.

With the differences in the number of available predictions for each VEP, these metrics are not straightforward to use to compare methods directly. Therefore, the rank score, that is based only on shared variant predictions, seems better suited for a fair comparison between VEPs.

The best VEPs based on the rank score are still MetaRNN, BayesDel_addAF and ClinPred with individual rank scores of 0.938, 0.938, and 0.898 respectively (see Figure 2C and Table 2). A rank score of 0.938 for MetaRNN means that MetaRNN significantly beats 93% of evaluated VEPs on the ClinVar dataset using shared variants. The rank score can change the rankings of VEPs that may appear to be very similar in performance using classical metrics. For example, EVE and DANN have close AUROC values with 0.896 and 0.890 respectively. With the rank scores, the distributions are reversed and EVE highly outperforms DANN with a rank score of 0.719 compared to 0.344 for DANN. Similarly, Eigen is ahead of VEPs like SIFT, PolyPhen-2 on AUROC metric (see Figure S3A), but using rank scores metric, it appears that Eigen performances are not significantly better than cited VEPs, and is actually below them when compared using shared variants (see Figure 2C).

This illustrates how rank score can alter the perceived performance among VEPs, highlighting its importance in such evaluation. Nevertheless, it must be noticed that the rank score of EVE must be considered with caution since the amount of data used to compute this rank score is very low and might not be representative of EVE true performance.

##### 2.3.1.2. Performances on high quality variants from ClinVar

ClinVar_HQ is a subset of the initial ClinVar dataset consisting of high quality variants selected as described in section 1.1.1. ClinVar_HQ dataset provided slightly lower performances with an AUROC median value of 0.88 against 0.93 for the ClinVar dataset (see Figure 2A and Table 3). MCC values shared the same tendency with a median MCC of 0.61 for ClinVar versus 0.59 for ClinVar_HQ (see Figure 2B). The top performing VEPs remain MetaRNN, BayesDel and ClinPred using the AUROC or MCC metrics (see Figure S3B and S4B). Interestingly, using rank score metric, these VEPs are barely in the top 5 of best VEPs, except for MetaRNN. Indeed, AlphaMissense reached the first position with a rank score of 0.766 followed by MetaRNN with a score of 0.734 (see Figure 2D), tied with SIGMA. These performances are interesting as these variants have better annotation quality; nevertheless they are also associated with a limited number of occurrences.

**Table 3.**
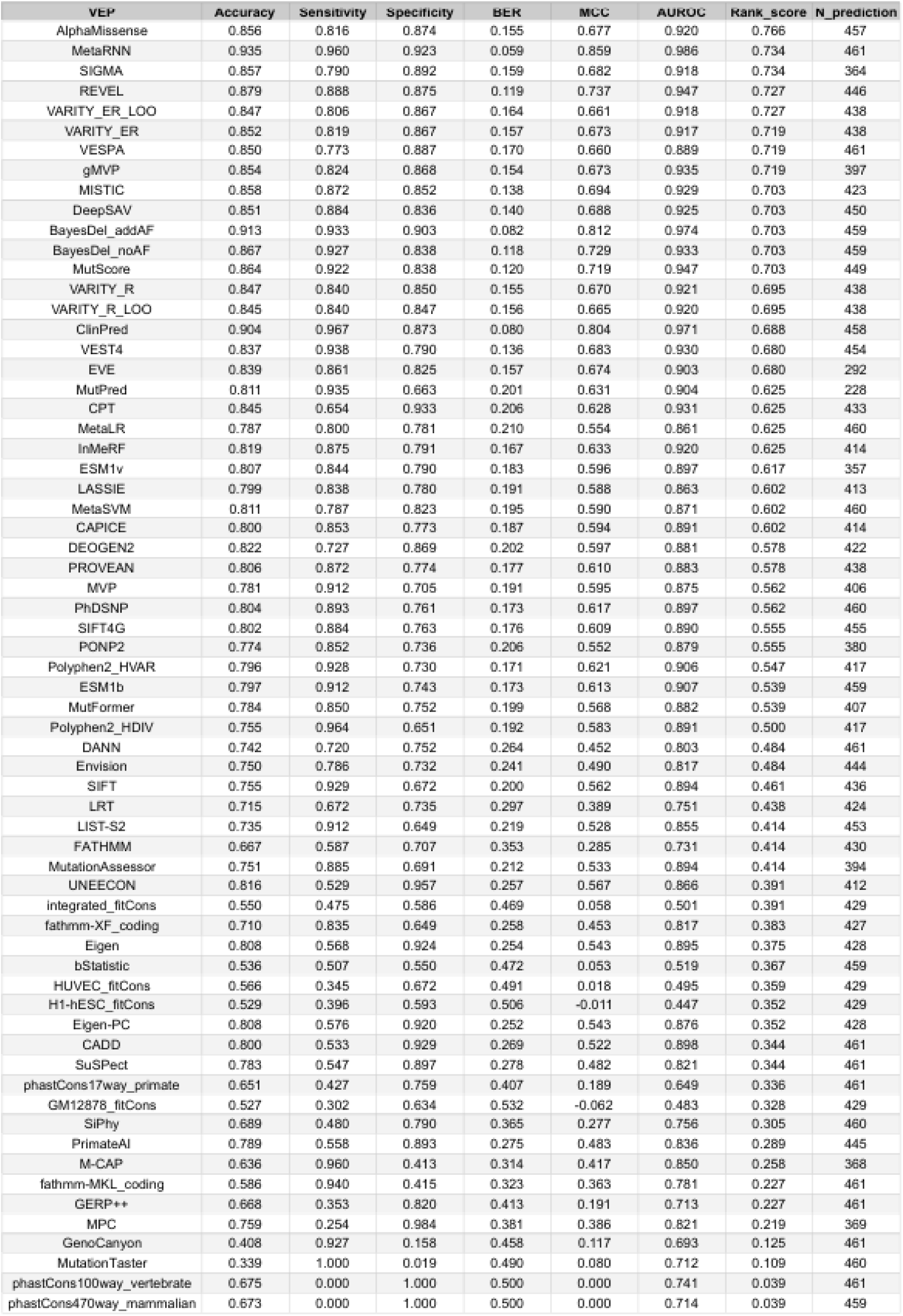
ClinVar_HQ performances. Overview of performances of each VEP on ClinVar_HQ dataset. Rank score and the number of predictions available are also provided.

#### 2.3.2. Clinical variants

On the Clinical dataset, performances are clearly decreasing compared to the ClinVar results. The median AUROC value is 0.73 (see Figure 3A) versus 0.93 for ClinVar (see Figure 2A). Similarly, the median MCC falls sharply to 0.32 (see Figure 3B), compared with 0.61 for ClinVar (see Figure 2B).

**Figure 3.**
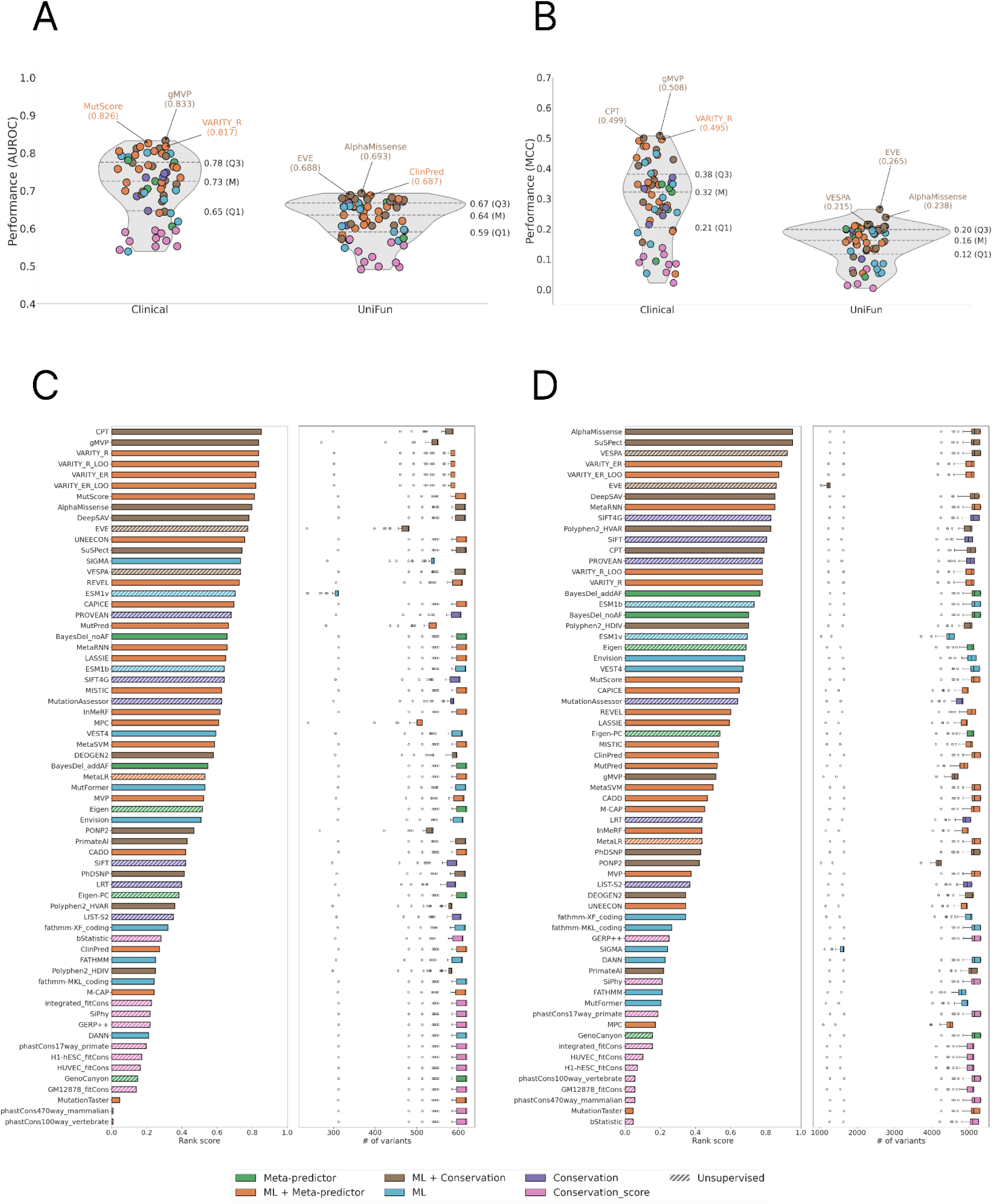
Rank score performances in Clinical and UniFun datasets. (above) Distribution of AUROC (A) and MCC (B) values for Clinical and UniFun datasets. (below) Histogram of rank score values for Clinical (C) and UniFun (D) datasets. see Figure 2 for additional information. Visible outliers correspond to comparisons that involve VEPs that have low data coverage.

Still numerous meta-methods are found in the top performing VEPs like VARITY (all four versions) or MutScore (see Table 4). However, ClinPred and BayesDel, which are meta-methods and were among the top performing VEPs on the ClinVar dataset, are now outside the top10 (see Figures S3C and S4C). Rank score analysis shows that CPT, gMVP and all four versions of VARITY are dominating this dataset (see Figure 3C) with rank scores ranging from 0.82 to 0.852. These VEPs are followed by MutScore, AlphaMissense and DeepSAV that have close rank scores. Globally, VEP’s predictions are less accurate on this dataset composed of clinical variants compared to ClinVar dataset. It appears that this specific and small dataset, composed of manually examined clinical variants, is more challenging for all the VEPs considered. In parallel, some VEPs like VARITY, MutScore and AlphaMissense show consistency on being in the top performing VEPs in ClinVar, ClinVar_HQ and in the Clinical dataset (see Figures 2 and 3).

**Table 4.**
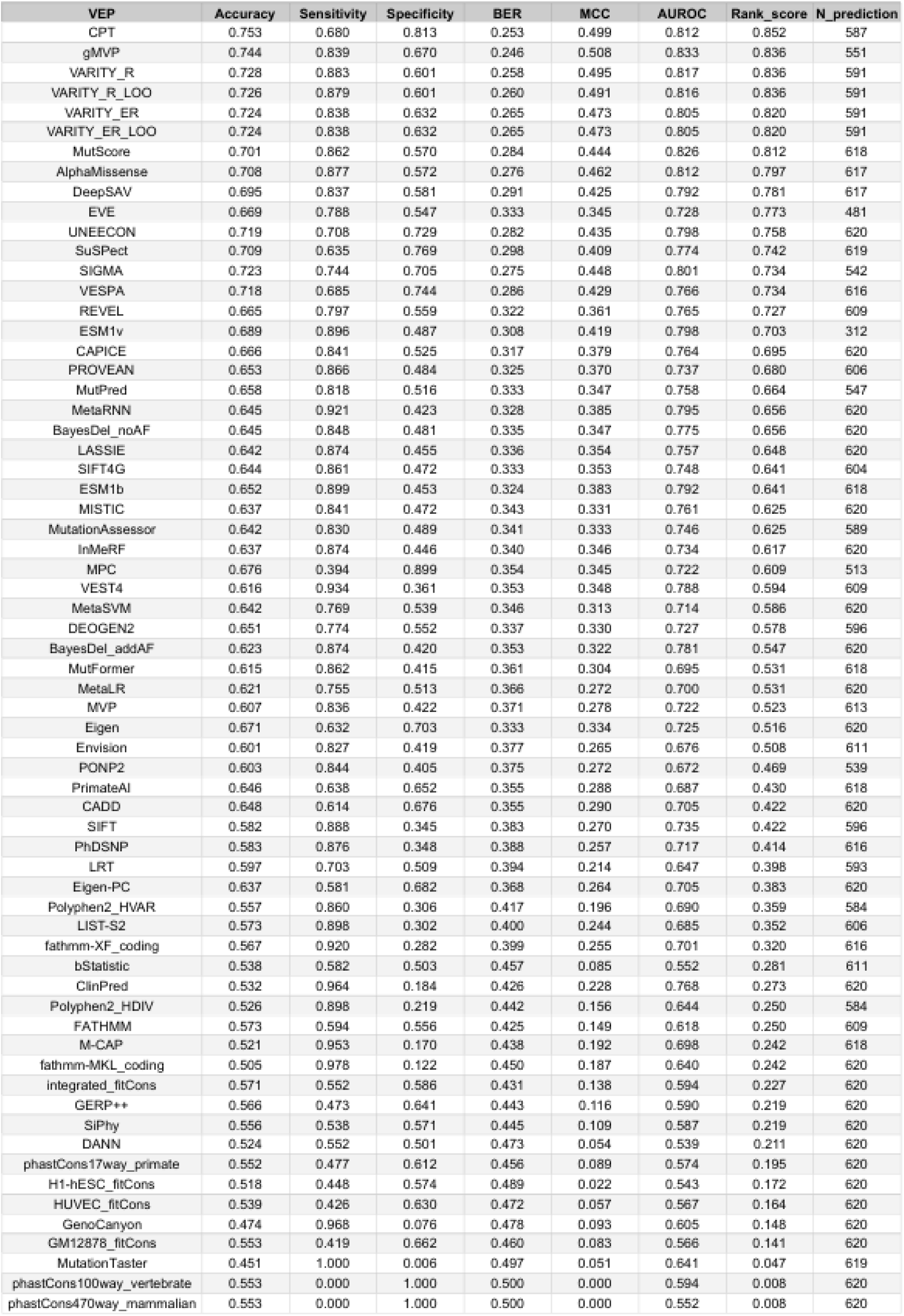
Clinical performances. Overview of performances of each VEP on the Clinical dataset. Rank score and the number of predictions available are also provided.

| VEP | Accuracy | Sensitivity | Specificity | BER | MCC | AUROC | Rank_score | N_prediction |
| --- | --- | --- | --- | --- | --- | --- | --- | --- |
| CPT | 0.753 | 0.680 | 0.813 | 0.253 | 0.499 | 0.812 | 0.852 | 587 |
| gMVP | 0.744 | 0.839 | 0.670 | 0.246 | 0.508 | 0.833 | 0.836 | 551 |
| VARITY_R | 0.728 | 0.883 | 0.601 | 0.258 | 0.495 | 0.817 | 0.836 | 591 |
| VARITY_R_LOO | 0.726 | 0.879 | 0.601 | 0.260 | 0.491 | 0.816 | 0.836 | 591 |
| VARITY_ER | 0.724 | 0.838 | 0.632 | 0.265 | 0.473 | 0.805 | 0.820 | 591 |
| VARITY_ER_LOO | 0.724 | 0.838 | 0.632 | 0.265 | 0.473 | 0.805 | 0.820 | 591 |
| MutScore | 0.701 | 0.862 | 0.570 | 0.284 | 0.444 | 0.826 | 0.812 | 618 |
| AlphaMissense | 0.708 | 0.877 | 0.572 | 0.276 | 0.462 | 0.812 | 0.797 | 617 |
| DeepSAV | 0.695 | 0.837 | 0.581 | 0.291 | 0.425 | 0.792 | 0.781 | 617 |
| EVE | 0.669 | 0.788 | 0.547 | 0.333 | 0.345 | 0.728 | 0.773 | 481 |
| UNEECON | 0.719 | 0.708 | 0.729 | 0.282 | 0.435 | 0.798 | 0.758 | 620 |
| SuSPect | 0.709 | 0.635 | 0.769 | 0.298 | 0.409 | 0.774 | 0.742 | 619 |
| SIGMA | 0.723 | 0.744 | 0.705 | 0.275 | 0.448 | 0.801 | 0.734 | 542 |
| VESPA | 0.718 | 0.685 | 0.744 | 0.286 | 0.429 | 0.766 | 0.734 | 616 |
| REVEL | 0.665 | 0.797 | 0.559 | 0.322 | 0.361 | 0.765 | 0.727 | 609 |
| ESM1v | 0.689 | 0.896 | 0.487 | 0.308 | 0.419 | 0.798 | 0.703 | 312 |
| CAPICE | 0.666 | 0.841 | 0.525 | 0.317 | 0.379 | 0.764 | 0.695 | 620 |
| PROVEAN | 0.653 | 0.866 | 0.484 | 0.325 | 0.370 | 0.737 | 0.680 | 606 |
| MutPred | 0.658 | 0.818 | 0.516 | 0.333 | 0.347 | 0.758 | 0.664 | 547 |
| MetaRNN | 0.645 | 0.921 | 0.423 | 0.328 | 0.385 | 0.795 | 0.656 | 620 |
| BayesDel_noAF | 0.645 | 0.848 | 0.481 | 0.335 | 0.347 | 0.775 | 0.656 | 620 |
| LASSIE | 0.642 | 0.874 | 0.455 | 0.336 | 0.354 | 0.757 | 0.648 | 620 |
| SIFT4G | 0.644 | 0.861 | 0.472 | 0.333 | 0.353 | 0.748 | 0.641 | 604 |
| ESM1b | 0.652 | 0.899 | 0.453 | 0.324 | 0.383 | 0.792 | 0.641 | 618 |
| MISTIC | 0.637 | 0.841 | 0.472 | 0.343 | 0.331 | 0.761 | 0.625 | 620 |
| MutationAssessor | 0.642 | 0.830 | 0.489 | 0.341 | 0.333 | 0.746 | 0.625 | 589 |
| InMeRF | 0.637 | 0.874 | 0.446 | 0.340 | 0.346 | 0.734 | 0.617 | 620 |
| MPC | 0.676 | 0.394 | 0.899 | 0.354 | 0.345 | 0.722 | 0.609 | 513 |
| VEST4 | 0.616 | 0.934 | 0.361 | 0.353 | 0.348 | 0.788 | 0.594 | 609 |
| MetaSVM | 0.642 | 0.769 | 0.539 | 0.346 | 0.313 | 0.714 | 0.586 | 620 |
| DEOGEN2 | 0.651 | 0.774 | 0.552 | 0.337 | 0.330 | 0.727 | 0.578 | 596 |
| BayesDel_addAF | 0.623 | 0.874 | 0.420 | 0.353 | 0.322 | 0.781 | 0.547 | 620 |
| MutFormer | 0.615 | 0.862 | 0.415 | 0.361 | 0.304 | 0.695 | 0.531 | 618 |
| MetaLR | 0.621 | 0.755 | 0.513 | 0.366 | 0.272 | 0.700 | 0.531 | 620 |
| MVP | 0.607 | 0.836 | 0.422 | 0.371 | 0.278 | 0.722 | 0.523 | 613 |
| Eigen | 0.671 | 0.632 | 0.703 | 0.333 | 0.334 | 0.725 | 0.516 | 620 |
| Envision | 0.601 | 0.827 | 0.419 | 0.377 | 0.265 | 0.676 | 0.508 | 611 |
| PONP2 | 0.603 | 0.844 | 0.405 | 0.375 | 0.272 | 0.672 | 0.469 | 539 |
| PrimateAI | 0.646 | 0.638 | 0.652 | 0.355 | 0.288 | 0.687 | 0.430 | 618 |
| CADD | 0.648 | 0.614 | 0.676 | 0.355 | 0.290 | 0.705 | 0.422 | 620 |
| SIFT | 0.582 | 0.888 | 0.345 | 0.383 | 0.270 | 0.735 | 0.422 | 596 |
| PhDSNP | 0.583 | 0.876 | 0.348 | 0.388 | 0.257 | 0.717 | 0.414 | 616 |
| LRT | 0.597 | 0.703 | 0.509 | 0.394 | 0.214 | 0.647 | 0.398 | 593 |
| Eigen-PC | 0.637 | 0.581 | 0.682 | 0.368 | 0.264 | 0.705 | 0.383 | 620 |
| Polyphen2_HVAR | 0.557 | 0.860 | 0.306 | 0.417 | 0.196 | 0.690 | 0.359 | 584 |
| LIST-S2 | 0.573 | 0.898 | 0.302 | 0.400 | 0.244 | 0.685 | 0.352 | 606 |
| fathmm-XF_coding | 0.567 | 0.920 | 0.282 | 0.399 | 0.255 | 0.701 | 0.320 | 616 |
| bStatistic | 0.538 | 0.582 | 0.503 | 0.457 | 0.085 | 0.552 | 0.281 | 611 |
| ClinPred | 0.532 | 0.964 | 0.184 | 0.426 | 0.228 | 0.768 | 0.273 | 620 |
| Polyphen2_HDIV | 0.526 | 0.898 | 0.219 | 0.442 | 0.156 | 0.644 | 0.250 | 584 |
| FATHMM | 0.573 | 0.594 | 0.556 | 0.425 | 0.149 | 0.618 | 0.250 | 609 |
| M-CAP | 0.521 | 0.953 | 0.170 | 0.438 | 0.192 | 0.698 | 0.242 | 618 |
| fathmm-MKL_coding | 0.505 | 0.978 | 0.122 | 0.450 | 0.187 | 0.640 | 0.242 | 620 |
| integrated_fitCons | 0.571 | 0.552 | 0.586 | 0.431 | 0.138 | 0.594 | 0.227 | 620 |
| GERP++ | 0.566 | 0.473 | 0.641 | 0.443 | 0.116 | 0.590 | 0.219 | 620 |
| SiPhy | 0.556 | 0.538 | 0.571 | 0.445 | 0.109 | 0.587 | 0.219 | 620 |
| DANN | 0.524 | 0.552 | 0.501 | 0.473 | 0.054 | 0.539 | 0.211 | 620 |
| phastCons17way_primate | 0.552 | 0.477 | 0.612 | 0.456 | 0.089 | 0.574 | 0.195 | 620 |
| H1-hESC_fitCons | 0.518 | 0.448 | 0.574 | 0.489 | 0.022 | 0.543 | 0.172 | 620 |
| HUVEC_fitCons | 0.539 | 0.426 | 0.630 | 0.472 | 0.057 | 0.567 | 0.164 | 620 |
| GenoCanyon | 0.474 | 0.968 | 0.076 | 0.478 | 0.093 | 0.605 | 0.148 | 620 |
| GM12878_fitCons | 0.553 | 0.419 | 0.662 | 0.460 | 0.083 | 0.566 | 0.141 | 620 |
| MutationTaster | 0.451 | 1.000 | 0.006 | 0.497 | 0.051 | 0.641 | 0.047 | 619 |
| phastCons100way_vertebrate | 0.553 | 0.000 | 1.000 | 0.500 | 0.000 | 0.594 | 0.008 | 620 |
| phastCons470way_mammalian | 0.553 | 0.000 | 1.000 | 0.500 | 0.000 | 0.552 | 0.008 | 620 |

#### 2.3.3. Functional variants

The UniFun dataset is particularly different from the two previous datasets. Indeed, UniFun is composed of variants that impact (or not) the function of the protein, verified experimentally through mutagenesis experiment, but with no direct link with the clinical aspect of the variant. Interestingly, performances on this dataset are highly different from the two previous datasets. Indeed, AUROC (see Figure 3A) and MCC (see Figure 3B) medians are strongly decreasing, with a median of 0.64 for AUROC and only 0.16 for MCC. Those values are quite low compared to the performances on ClinVar or Clinical datasets. Considering the AUROC metric, the best performing VEPs on this dataset are AlphaMissense, EVE and ClinPred with close AUROC values of 0.693, 0.688 and 0.687 respectively (see Figure S3D). However, EVE surpasses other top VEPs regarding the MCC metric with a MCC of 0.265, ahead of AlphaMissense (0.238) and VESPA (0.215) (see Figures 3B and S4D, and Table 5).

**Table 5.**
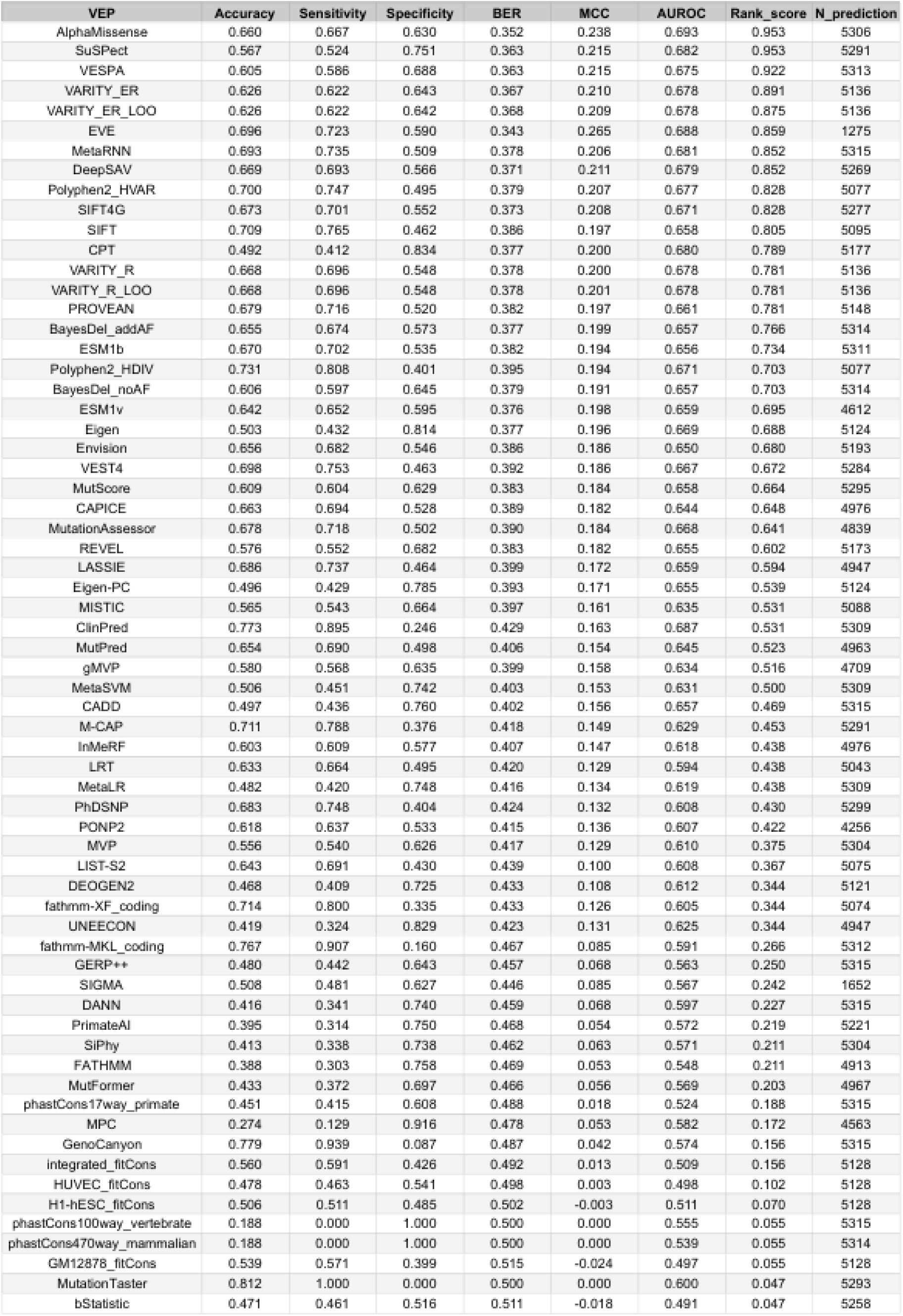
UniFun performances. Overview of performances of each VEP on the UniFun dataset. Rank score and the number of predictions available are also provided.

Using rank score analysis, the ranks of VEPs slightly change (see Figure 3D). AlphaMissense reached the first position with a rank score of 0.953 tied with SuSPect and followed by VESPA having a rank score of 0.922, VARITY at 0.891 and 0.875 for both ER and ER_LOO versions, and EVE at 0.859 (see Table 5).

We can notice that SIFT4G is reaching the top 10 with a rank score of 0.828 (and 0.805 for SIFT), indicating that even ancient algorithms are able to compete with recent ones on this dataset. Overall, this dataset is the most difficult for VEPs, as their predictive values are all quite low. It is also reasonable given the absence of a simple link with the pathology question.

### 2.4. Performances overview using other metrics

#### 2.4.1. Sensitivity and specificity insights

As shown in the previous sections, it is crucial to apply multiple measurements to evaluate the different methods with the datasets. Sensitivity and specificity are commonly used metrics to assess the performances of VEP. Respectively, they inform about the ability of the VEP to correctly predict the pathogenic (or positive label) and benign (or negative label) and therefore detect a potential bias in favor of one specific class. Sensitivity values are quite similar with a median value of 0.80, 0.84 for respectively ClinVar and Clinical datasets, while UniFun reaches only 0.62 (see Figure S5A). VEPs are fairly performing in predicting the pathogenic label. They appear to be slightly better predicted in the Clinical than ClinVar dataset and more poorly predicted in the UniFun dataset. Nonetheless, the specificity values are very high in the ClinVar dataset, meaning that benign variants are very well predicted (see Figure S5B). On the Clinical and UniFun datasets, specificity values are quite low and more sparse, indicating that VEPs struggle to correctly predict benign labels on these datasets. Some VEPs are among the best in sensitivity like MutationTaster that reaches a sensitivity of almost 1 for all datasets, but also a specificity close to 0 (see Tables 2, 3 and 4). This imbalance indicates bias toward the pathogenic label and more particularly that MutationTaster predicts every label as pathogenic. This can also be detected with the BER analysis, where such VEP will have a high BER value (see Figure S5C). A VEP that has a good balance between predicting benign and pathogenic labels will likely have a low BER. For the ClinVar dataset, ClinPred, MetaRNN and BayesDel_addAF have the lowest BER with very low values (around 0). On the Clinical dataset, BER values are higher than in ClinVar with gMVP, CPT and VARITY_R being the best VEPs with BER values around 0.25. Finally, on the UniFun dataset, BER values are even higher ranging from 0.343 to 0.363 with EVE, AlphaMissense and VESPA respectively. MCC and BER metrics are consistent with each other by yielding the same top three ranking of VEPs for each dataset (see Figures 2B, 3B and S5C).

Overall, the ClinVar dataset seems to incorporate more benign variants that are easy to predict which may inflate measured performances, reflected by very high specificity values (see Figure S5B). On the contrary, the Clinical and UniFun datasets are more balanced in terms of sensitivity and specificity.

#### 2.4.2. Average performance on clinical variants

One way to calculate a global performance over clinical variants is to compute the mean rank score of rank scores previously computed on ClinVar and Clinical datasets. This choice is motivated by the high representativeness of the rank score to estimate VEP’s performance in contrast to other ones, which can not easily be done with other metrics. We used ClinVar_HQ dataset instead of ClinVar to compute the mean rank score, because of the higher reliability of this dataset compared to the initial ClinVar dataset. By doing the mean of rank scores from ClinVar_HQ and Clinical datasets, we found that the best performing VEP overall is AlphaMissense with a mean rank score of 0.781 (see Figure 4), meaning that on average, AlphaMissense significantly beats 78% of other tested VEPs on ClinVar_HQ (i.e. clinical variants from ClinVar) and Clinical datasets. The next best VEPs are gMVP having a mean rank score of 0.777 and VARITY_ER_LOO with a mean rank score of 0.773. These VEPs are followed by remaining versions of VARITY and MutScore with rank scores ranging from 0.758 to 0.77, representing the best performing meta-methods. Next VEPs, such as DeepSAV and CPT, belong to the ML + Conservation group (like AlphaMissense) and have mean rank scores of 0.742 and 0.738 respectively. SIGMA (the only one in the ML group) has a mean rank score of 0.734, followed by VESPA, EVE and REVEL have the same average rank scores 0.727. All of these VEPs are the ones that have a mean rank score above 0.7 and constitute the group of VEPs of “Highest performances”. VEPs with a mean rank score that are between 0.7 and 0.6 are grouped in the “Good performances” group and this include VEPs such as MetaRNN, BayesDel, MISTIC or PROVEAN. The third group, called “Medium performances”, is composed of VEPs with mean rank scores between 0.6 and 0.5, such as PONP2 or ESM1b. The “Low performances” group is composed of VEPs having mean rank scores between 0.5 and 0.3 such as CADD, PrimateAI or DANN. Finally, the “Lowest performances” group is composed of remaining VEPs having a mean rank score below 0.3. Most conservation scores, such as fitCons, phastCons or GERP++ are in this group but also GenoCanyon or MutationTaster which are meta-methods.

**Figure 4.**
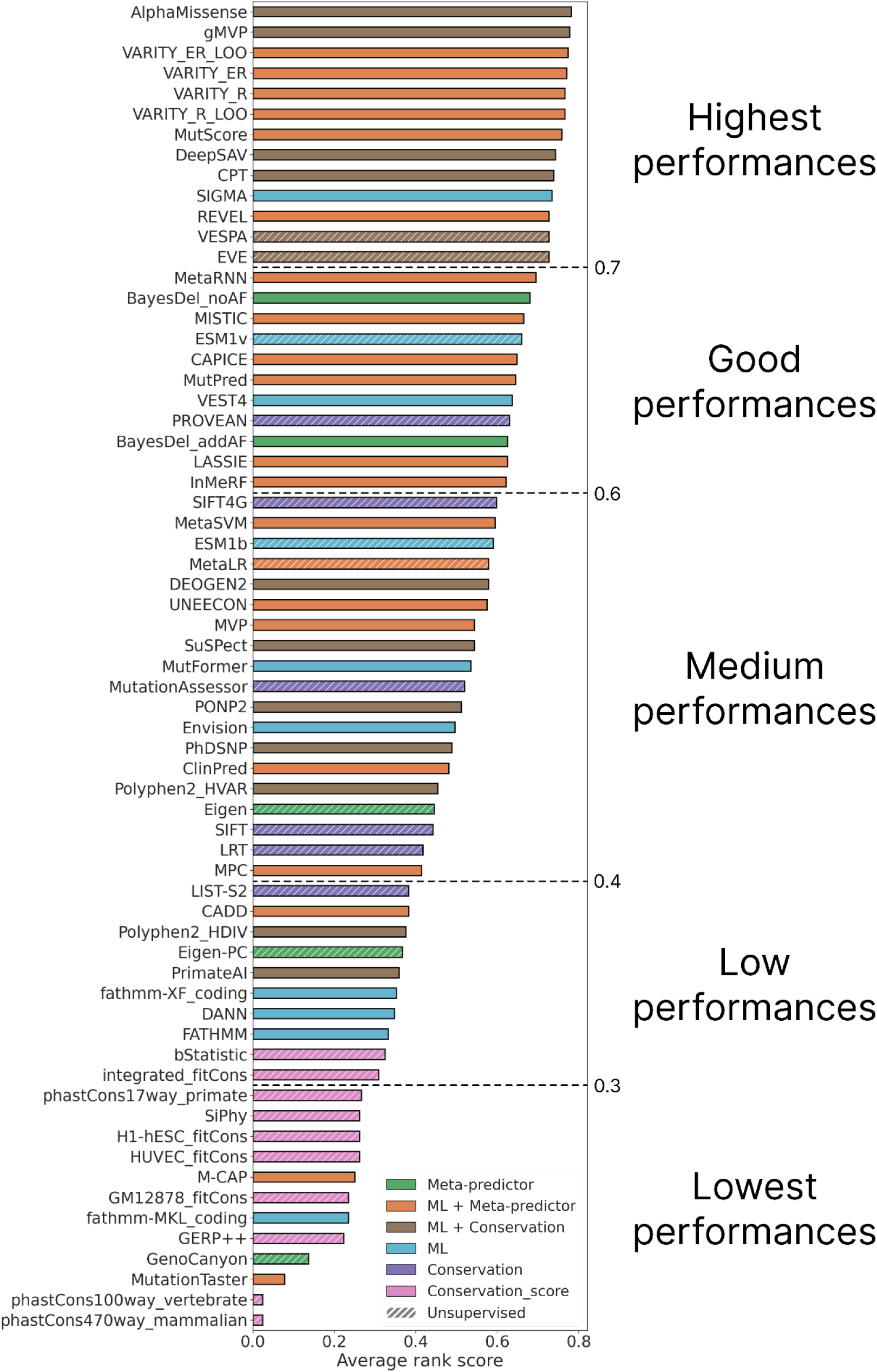
Average rank score and group of performance for clinical variants. The average rank score represents the mean between rank scores values from ClinVar_HQ and Clinical datasets and constitute the global performance of each VEP on clinical variants. Different cutoff values are used to regroup VEPs according to their performance, i.e. 0.7, 0.6, 0.4 and 0.3, resp.

Addition of UniFun dataset to the analysis (see Figure S6) slightly modify the results, but keep AlphaMisense as the most efficient approach. Most approaches remain within their performance classes; however, some VEPs achieve higher ranks, such as MetaRNN which moves into the highest performance group with a mean rank score of 0.734.

Incorporating multiple VEPs from various knowledge sources appears to be a good and efficient way to build a VEP that performs decently on various variant contexts as shown by VARITY, MetaRNN and MutScore. However, AlphaMissense demonstrates that non meta-methods can also perform very well and sometimes with more consistency than meta-methods. Globally, evolutionary information appears to be essential to get the best performances in predicting impact of variants as numerous top performing VEPs are using MSA derived information.

#### 2.4.3. Performances for each VEP type

Some VEP types may be more suited to specific types of variants. Variants from ClinVar seem to be better predicted by meta-methods (see Figure S7A) with better rank scores for meta-methods associated with machine learning algorithms (ML + Meta-predictor). The ClinVar_HQ dataset shows that the difference between meta-methods with machine learning (ML + Meta-predictor group) and VEP using conservation information with machine learning (ML + Conservation group) is not significant, as they achieve similar results on this dataset (see Figure S7B). These tendencies are also found for the Clinical dataset (see Figure S7C). However, for the UniFun dataset, it is clear that VEPs using conservation information clearly outperform, whether they use machine learning algorithms or not (Conservation and ML + Conservation groups, see Figure S7D). These results suggest that the type of variant analyzed (functional or pathogenic) may help to decide which VEP type is more appropriate to use. For clinical variants, meta-methods using machine learning seem to be better suited (*e.g.*, MetaRNN, VARITY or MutScore), whereas for functional variants, VEPs using conservation information are a better choice with a preference for those using machine learning (*e.g.*, AlphaMissense, SuSPect, VESPA or EVE).

### 2.5. Variant difficulty

Thus, VEPs are performing well on the ClinVar dataset, but struggle on the Clinical and even more on the UniFun dataset. These results raise the question of why the variants present in these two datasets are more difficult to predict. We analyze our results in more depth to classify the variations according to their level of difficulty, as this could be linked to specific characteristics. An error rate per variant was built, ranging from 0 to 1, corresponding to the proportion of wrong predictions amongst all VEP’s predictions for each variant. Figure 5A shows the distribution of error rates for benign and pathogenic variants. Based on these observations, we propose to classify the prediction variant difficulty in three classes: “easy” variants concern variants with an error rate less than 30%, “hard” variants with an error rate superior to 70%, remaining variants being classified into the “moderate” class.

**Figure 5.**
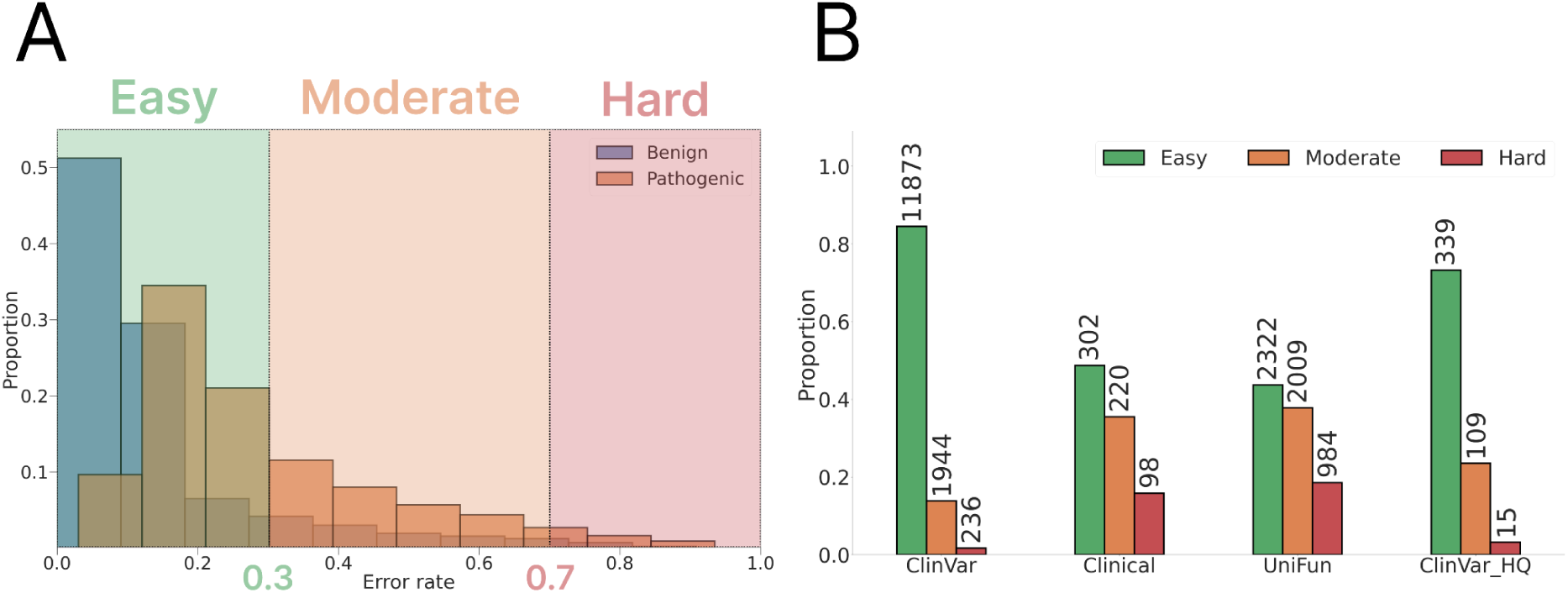
Determination of variant difficulty classes. (A) Example of distribution of error rates in ClinVar dataset and determination of easy, moderate and hard classes. easy variants have an error rate below 20%, hard variants have an error rate above 70%, remaining variants are classified as moderate variants. (B) Overview of variant difficulty classes proportion in each dataset.

As expected by the high performances of our pool of VEPs, ClinVar has the greatest proportion of easy variants with 11,873 variants (84.4%), and the least proportion of hard variants with 236 variants (1.6%), the moderate class corresponds to 1944 variants (remaining 13.7%) (see Figure 5B). However, Clinical and UniFun datasets have respectively 49% and 43% of easy variants, i.e. is half of ClinVar proportion, and 15% and 18% of hard variants, which is ten times more than in ClinVar. hard variants are likely challenging VEPs by dropping down their performances. In the following sections, we tried to identify some biological characteristics that could be associated with the prediction difficulty.

#### 2.5.1. Amino acid composition

The amino acid composition of the three classes were analyzed. Amino acids were clustered according to their physicochemical group among: aromatic, negatively and positively charged, polar and nonpolar. No specific tendencies were observed for the wild-type residue (see Figures S8A, S8C and S8E) or the mutated residue (see Figures S8B, S8D and S8F) toward each class. These results show that sequence information alone is not sufficient to biologically characterize easy or hard variants.

#### 2.5.2. Solvent accessibility

To look at each protein of our datasets, AlphaFold2 structural models and experimental structures from the PDB were retrieved in order to analyze the solvent accessibility surface area for each variant. For AlphaFold2 models, only variant positions that are modeled with pLDDT values higher than 80 are kept, *i.e.* sufficient to consider that the model is of correct quality for this position.

Amino acid variation could create a difference in the interacting network within the protein itself. Sometimes, the modified network could impact the protein so deeply that its proper integrity could be altered, leading to a possible strong and probably negative impact on the function, stability and structure of the protein. From an average perspective, those mutations happen in the core of the protein, a region that is crucial for its function. A mutation located at the protein surface has less chance to create a very unstable interaction network and is less probable to be deleterious.

Solvent accessibility analysis shows that easy variants correspond to these two cases described. easy benign variants are indeed located at the surface of the protein whereas easy pathogenic variants are located in the core of the protein (see Figure 6). This result is found in both filtered AlphaFold2 structures and also for experimental structures.

**Figure 6.**
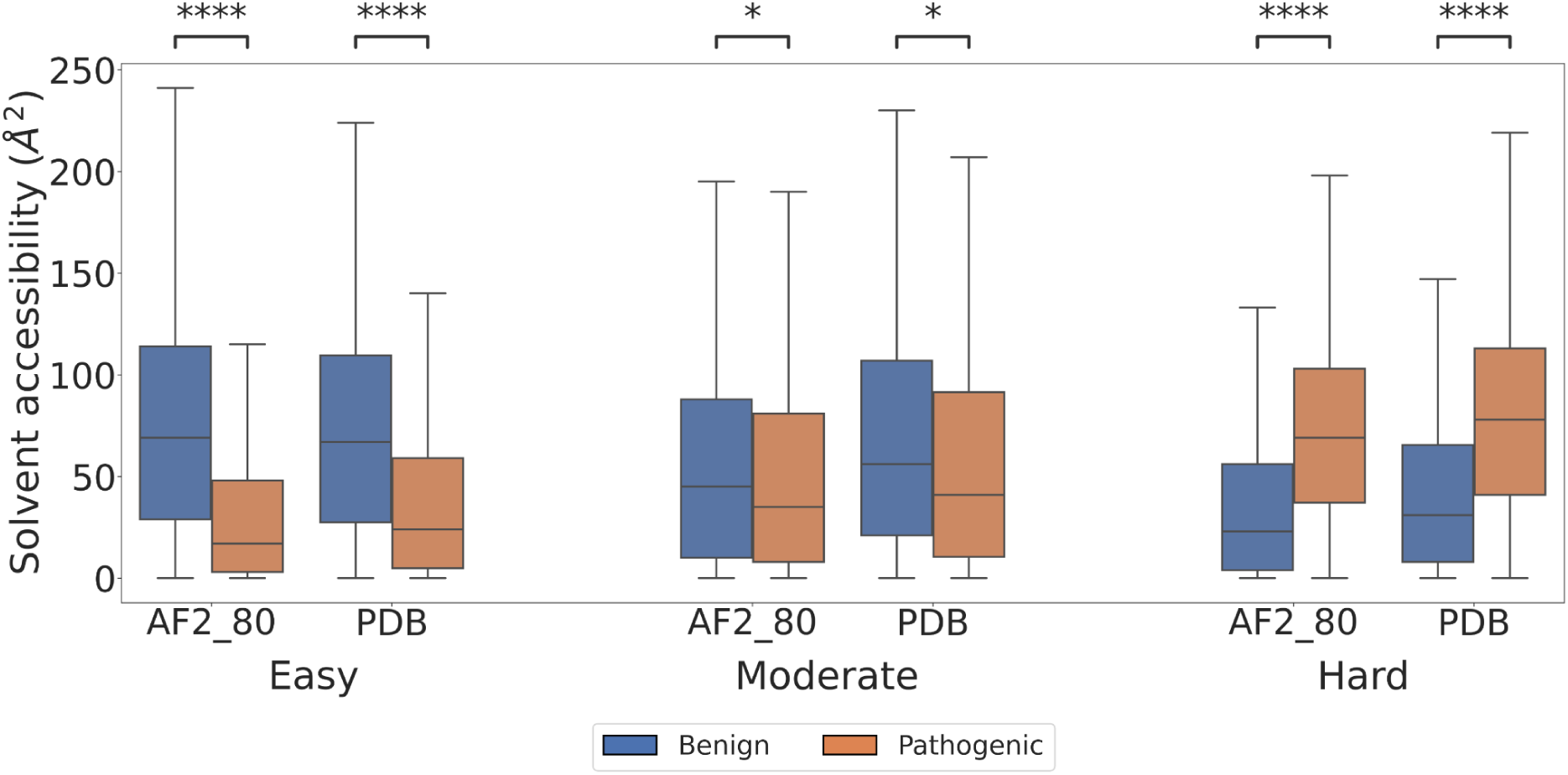
Solvent accessibility analysis on concatenated data. Solvent accessibility values for easy, moderate and hard variants. Here, all three datasets have been concatenated. Solvent accessibility values for benign variants are represented in blue and for pathogenic variants in orange using protein structure from the AF2 database filtered at 80 pLDDT (AF2_80, see methods) or from the PDB. Mann-Whitney statistical test has been applied for each comparison.

However, it is totally reversed for hard variants. Indeed, hard benign variants are mostly buried within the protein structure while hard pathogenic variants are located at the surface, exposed to solvent. These results suggest that actual VEPs have learned that the variant labeling is correlated to the residue solvent exposure following a simple pattern: a benign variant is located at the surface and a pathogenic variant is buried by preference. Nevertheless, there are still some variants that are misinterpreted by VEPs, probably because they do not correspond to a general scheme of the classical variant.

To ensure that this observed signal is not influenced by any particular dataset that may have a disproportionate impact on the analysis, we conducted the same analysis but with separated datasets (see Figure S9). We observed this tendency in both the ClinVar and UniFun datasets, with statistically significant differences (see Figures S9A and S9C). We also have a similar trend in the smaller Clinical dataset (see Figure S9B and Table 6), although the differences were not statistically significant due to the low number of data. This suggests that this categorization into “easy”, “moderate” and “hard” class may be extrapolated to variants whether it is a clinical or a functional variant.

**Table 6.** Amount of data used for SASA analysis. For each dataset and each difficulty, the amount of variants available for either AF2_80 or PDB group are represented. The last line corresponds to the concatenation of all three datasets. Bold values correspond to cases where statistical tests could not be used.

|  |  | EASY |  | MODERATE |  | HARD |  |
| --- | --- | --- | --- | --- | --- | --- | --- |
|  |  | AF2_80 | PDB | AF2_80 | PDB | AF2_80 | PDB |
| ClinVar | Benign | 4210 | 372 | 646 | 104 | 118 | 21 |
|  | Pathogenic | 1176 | 210 | 382 | 76 | 34 | **5** |
| Clinical | Benign | 21 | **1** | 59 | 14 | 63 | **9** |
|  | Pathogenic | 162 | 27 | 40 | **8** | **3** | **0** |
| UniFun | Benign | 144 | 3 | 244 | 82 | 152 | 37 |
|  | Pathogenic | 1697 | 452 | 868 | 275 | 317 | 122 |
| Concatanated | Benign | 4375 | 376 | 949 | 200 | 333 | 67 |
|  | Pathogenic | 3035 | 689 | 1290 | 359 | 354 | 127 |

This new categorization can be illustrated using the αIIbβ3 integrin exemple. This integrin is responsible for the normal blood coagulation by binding ligands, such as fibrinogen, to form blood clots^88,89^. A malfunction of this protein may lead to an absence of blood coagulation, also known as Glanzmann Thrombasthenia (GT) disease^90^. It has been shown for several examples of GTs associated with the Calf-1 and Calf-2 domains of this complex that the variants in fact changed nothing at the level of the mutation, but had effects at longer distances, potentially allosteric effects, on loops of Calf-1 and Calf-2^91^. An important number of disease-causing variants have been reported in GT from literature ^92^ and most of them are present in ClinVar. Among GT variants in ClinVar, the mutation L214R is found to be an easy pathogenic variant to predict with an error rate of 23% (below 30%). The position 214 is completely buried (SASA of 0 Å²) into the domain called β-propeller, which is the headpiece of the αII subunit that carries the ligand-binding site^89^ (see Figure 7). This result is consistent with our previous SASA analysis, the wild type residue has 2 initial interactions with residue A225 (in orange in the upper sphere) whereas the variant has one new interaction (in red in the upper sphere) with residue Y174. This additional interaction may change internal stability of the domain and protein which potentially made this variant as pathogenic and thus, an easy case to predict.

**Figure 7:**
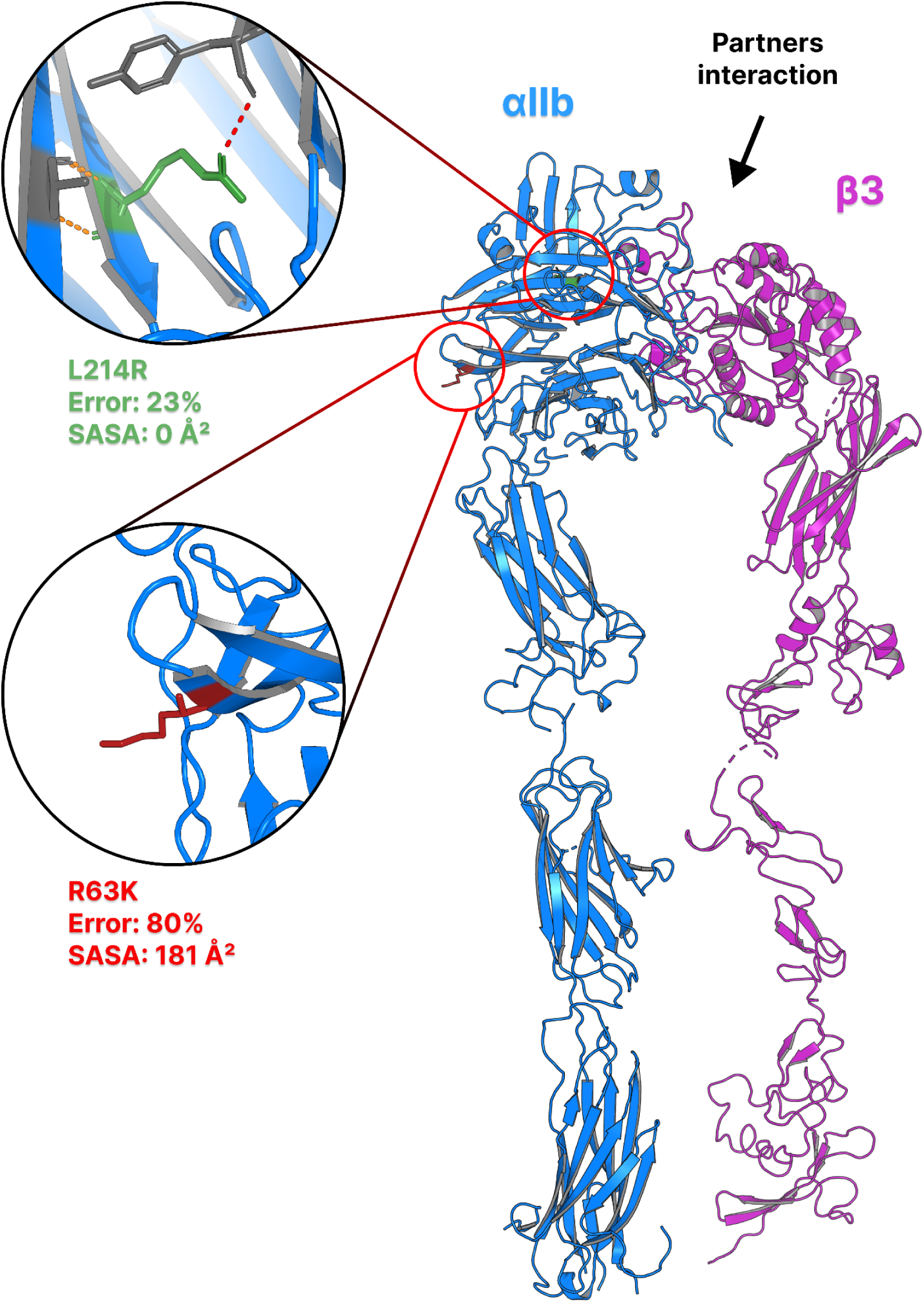
Example case of easy and hard pathogenic variants from integrin αIIbβ3. The extracellular domain of integrin αIIbβ3 is represented on the right using experimental structure from PDB 3FCS^94^. Originally in the PDB file, the integrin was in its bent or closed conformation, we opened it manually using PyMOL for representation purpose. On the left, two spheres represent specific variants on the αIIb subunit. The first sphere corresponds to the variant L214R (colored in green) with an error rate less than 30% and a SASA of 0 Å². The wildtype residue (L) is making two interactions with residue A225 (dashed lines in orange) while the mutant residue (R) has one additional interaction with residue Y174 (dashed line in red). The second sphere represents the variant R63K (colored in red) with an error rate of 80% and SASA of 181 Å².

Another pathogenic variant is the mutation R63K, present in ClinVar and in the same domain. This mutation has a very high error rate with 80% of VEPs failing to predict the correct annotation, placing it in the “hard” pathogenic group (error rate superior to 70%). This variant is less straightforward to interpret as it is located at the surface of the protein with a SASA of 181 Å² with no surrounding interaction. Moreover, the substitution does not indicate any strong physco-chemical change as the wild type (R) and mutant (K) residues are both positive and structurally very similar. Nonetheless, this variant is reported as pathogenic and has been deeply analyzed in the literature^93^. Authors of this study have found that the pathogenicity of this mutation is the result of a low level of mRNA, leading to a weak expression of the integrin at the surface of platelets. This reduced level of mRNA may be induced by other genomic events such as splicing as discussed in^93^. Such complex information is probably missing from VEP’s knowledge leading to incorrect pathogenicity prediction for most of them and high error rate.

#### 2.5.3. Function

We looked at whether certain protein functions were more likely to accumulate hard variants by retrieving protein function annotations from UniProt. All groups appear to be similarly populated with easy variants (see Figure 8) but some significant differences can be noticed (Table S1).

**Figure 8.**
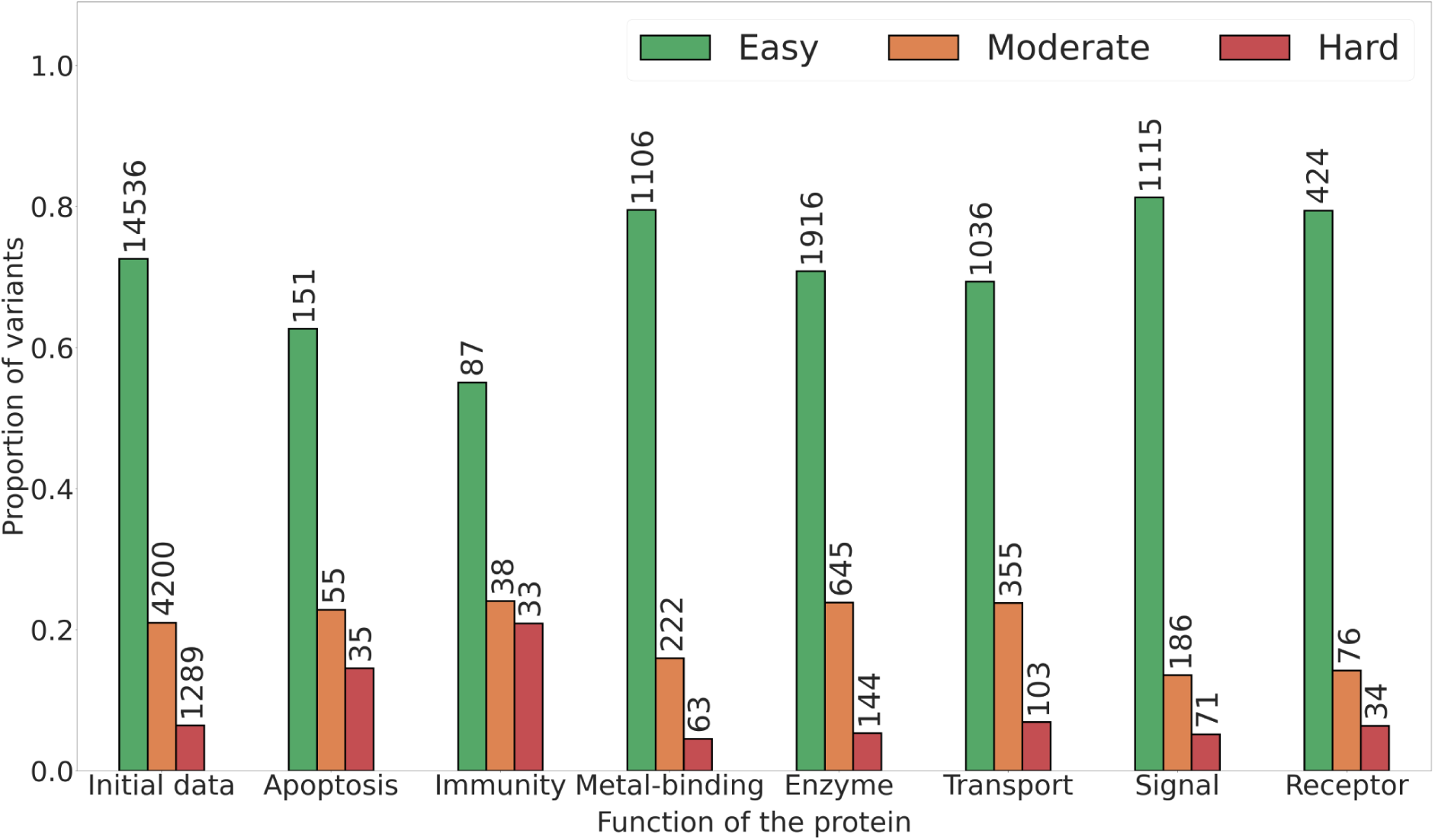
Overview of variant difficulty classes proportion across function. Amount of variants used to compute each proportion are represented above each bar for easy (green), moderate (orange) and hard (red) variants. Statistical tests have been computed between each pair and p-values are reported in Tables S1, S2 and S3.

Indeed, easy variants are significantly more detected in the “Signal” group than in “Immunity”, “Enzyme” or “Transport” groups (Tables S1 and S4). As for easy variants, we observed disparate statistical differences among moderate variants but with no clear tendency between protein functions. It appears that the “Signal” group has statistically less moderate variants than other groups (see Figure 8 and Tables S2 and S4).

“Apoptosis” and “Immunity” groups were significantly more populated with hard variants, which are 14.52% and 20.89% (Table S5). Remaining functions have between 4% and 6% of hard variants and are statistically less populated compared to “Apoptosis” and “Immunity” groups (Table S3). More interestingly, “Apoptosis” and “Immunity” groups are almost exclusively populated with pathogenic variants in the “hard” category (Table S4). This analysis reveals that hard variants are present in a differentially and statistically manner depending on the nature of protein function itself.

## Discussion

This benchmark analysis of 65 VEPs is an important key to understanding current advances and limitations of VEPs. Moreover, this study should help clinicians to better understand which tool they can use and why during their clinical routines.

The first analysis on ClinVar, the reference database with clinical classification of variants, shows that VEP’s performances are excellent despite our filtering steps to exclude variants that could lead to biased evaluation (see Figure 1).

On the ClinVar dataset, best performing VEPs are MetaRNN, ClinPred and BayesDel; these meta-methods are all trained on ClinVar. Despite our precautions (deletion of all ClinVar variants likely to have been learnt by these VEPs), we cannot eliminate the possibility that the VEPs have learnt signals specific to ClinVar variants (such as the type of genes), which could help to improve performance during evaluation. However, even unsupervised methods, such as VESPA or SIFT, have their performance inflated on ClinVar compared to Clinical or UniFun datasets despite having not learned any clinical label. It tends to indicate that the inflated performance is not related to label overfitting phenomena but rather to the nature of the dataset itself. Indeed, we have demonstrated that some variants are easier to predict than others. ClinVar is mainly composed of easy variants that are well predicted by most of VEPs. Benign variants were also better predicted compared to pathogenic ones, suggesting that these variants are easier to predict. Those results indicate that the database may contain some underlying bias that facilitates the inference of variant impact as suggested in^31^. Indeed, variants are annotated following the American College of Medical Genetics (ACMG) and Association for Molecular Pathology (AMP) classification^3^ that categorizes the variant either as a benign or pathogenic one. This tends to regroup variants that fit all these categories from the AMP classification, which ultimately favor the overrepresentation of concerned variants or genes. This overrepresentation can lead to an easier assessment of variant impact from ClinVar. Moreover, is it admitted that VEP tools can be used to help classify variants in ClinVar, meaning that the annotation determination may not be totally human based^3^. Therefore, evaluating or developing VEP using variants from ClinVar, partially annotated with the help of VEP tools, is kind of problematic and can bring new bias among ClinVar annotations for evaluation or training of VEPs. Moreover, this issue has been hypothesized as a new type 3 data circularity, described as an emerging bias induced by the huge and quick advances in VEP development as discussed in^31^. One way to prevent type 3 circularity could be to remove variants that have potentially been annotated with the help of VEP. Unfortunately, the use of VEP is not mentioned in ClinVar variant annotations.

Considering the ClinVar_HQ dataset, composed of variants with multiple submissions or have been verified manually by experts in ClinVar, performances are slightly lower compared to the initial ClinVar dataset but still very high. Top three VEPs on the initial ClinVar dataset are no longer there in ClinVar_HQ, leaving the first place to AlphaMissense tied with SuSPect. The quality selection on ClinVar variants has an impact on performances of VEPs. However, on both dataset, performance values are high and they both contain a significant and similar proportion of easy variants. By consequence, those results tend to affirm that the resulting “high quality” dataset seems to follow the same tendency, including also the potential pitfalls, as the initial ClinVar dataset. Even if one filters ClinVar variants to retain only variants of higher quality, an underlying bias is kept that facilitates the label prediction inflating performances.

Two other datasets were used in order to evaluate VEP using variants coming from different sources: Clinical and UniFun dataset. In the Clinical dataset, a high-quality dataset filtered from clinicians, we found that VEPs are much more challenged by these variants with lower AUROC and MCC values compared to performances on ClinVar datasets. The drop of performances can be explained by the sources of variants that are: (i) directly issued from clinical research, (ii) more filtered and (iii) are verified manually by experts. By consequence, the annotations and selected variants are of higher quality than the ClinVar dataset which can explain the important difficulty to make good predictions for VEPs. However we can notice that meta-methods are still among the best performing VEP.

The last one is the UniFun dataset which contains functional variants that are not clinical variants. We choose this dataset to analyze whether or not VEPs can still perform well on function variants. Overall, performance values are tremendously low with AUROC and MCC values not going further than 0.693 and 0.265, obtained by AlphaMissense and EVE respectively.

Except for VARITY and MetaRNN, all top VEPs are either from the group Conservation or in the Conservation + ML group (see Figure 3D). They all share similarities as they are using MSA in their algorithm. This indicates that evolutionary information extracted from MSA is crucial to predict the effect of functional variants. This seems consistent with the intrinsic nature of data itself, as those variants impact the function of the protein. Regions responsible for a protein function or its interaction ability with other proteins are generally well conserved across evolution and can be detected using MSA. Such information seems to be effectively extracted by VEPs using MSA in their algorithm and we show that they perform better on this dataset (see Figure S7D). It is surprising that even an ancient method such as SIFT can compete with the above-mentioned VEPs, indicating that even a simple method based on MSA information can extract enough evolutionary information to be competitive against more recent and technical tools in terms of prediction performance. VARITY and MetaRNN are performing also very well on this dataset despite the fact that they do not integrate explicitly conservation features. However, they are meta-methods that incorporate VEPs from the Conservation group such as SIFT, PROVEAN or LRT and conservation scores such as GERP++ or fitCons. These features can explain why VARITY and MetaRNN are performing so well on this special dataset. This highlights again the capacity of meta-methods to perform quite well thanks to the combination of multiple sources of knowledge about variant effects.

Considering average performances on clinical variants, we can regroup VEPs into 5 groups of performances as shown in Figure 4: (I) highest performances, (II) good performances, (III) medium performances, (IV) low performances and (V) lowest performances. Overall, the best performing VEP is AlphaMissense as it showed consistent performances on clinical variants (see Figure 4) with an average rank score of 0.781. It means that AlphaMissense is better than 78% of total VEPs on average on clinical variants but also across all datasets with UniFun (see Figure S6) on which it achieves an average rank score of 0.824. However, we should not exclude the usage of other high performing VEPs like VARITY, gMVP, MutScore, DeepSAV, CPT, VESPA, SIGMA, EVE or REVEL. They all have good average rank scores and are in the group “Highest performances” VEPs. MetaRNN also achieves to reach the high performance group when adding UniFun rank scores in the average computation (see Figure S6). Therefore, we recommend using these top-tier VEPs, in addition to AlphaMissense predictions, for a more representative assessment of variant impacts. These top-tier VEPs also have the advantage of being easily usable, since precomputed prediction files are available for all of them. Moreover, many of them are ready to use in the dbNSFP prediction database. EVE is also in the group of highest performing VEP but unfortunately, a very small amount of precomputed prediction is available for this tool, making it less easier to be used casually. Moreover, performance measures on EVE could be non representative of its true performances, because an important part of predictions were missing. Although this issue is partially solved with the rank score metric that evaluates VEPs using shared variants, the amount of missing data is so high that we cannot be sure about its overall rank. However, other benchmark analyses were done, analyzing EVE performances and showing that its performances were among the state of the art performances^30^, consistent with our results on limited variants available for EVE.

Conservation scores are among the lowest performing VEPs, suggesting that conservation scores used alone can not accurately assess the impact of variants. This is not surprising as they do not have been developed for this end but rather to detect evolutionary constraint regions of the gene. Despite this low performance, information extracted by conservation scores can be useful in combination with other sources of information to assess the impact of variants. Indeed, multiple meta predictors have incorporated conservation scores in their algorithm and have shown that they have non negligible importance in the final prediction through features importances analysis^28^.

One new insight gained by our benchmark is the difficulty of some variants to be correctly predicted. We have shown in our analysis that variants can be categorized into three groups of difficulties: (I) easy variants, (II) moderate variants and (III) hard variants. We found that these groups exhibit differences between them, indicating that considered variants probably have biological reasons on why they are easy and hard to predict. We show in our study that the solvent accessibility of the residues modified by the genetic variants can be one explanation that might classify the difficulty of the prediction. Indeed, easy variants correspond to ones that are located at the surface of the protein whereas pathogenic variants are located in the core of the protein. They follow a well-known scheme about the pathogenicity that can closely be linked to the inner protein function, which is usually buried into the protein. Hard variants correspond to the inverse situation, where benign variants are located in the core, and pathogenic variants at the surface of the protein.

These variants are ultimately hard to detect for VEPs and one potential reason for this challenge is the absence of specific input information about the impact of such variants. Indeed, mutations located in the protein core probably have a higher impact on its intrinsic function than one in the surface. However, this aspect is directly related to the nature of the protein function itself. Therefore, the variant difficulty was directly analyzed in regard to the function type of the protein coded by the genes listed in our three datasets. Hard pathogenic variants are in highest proportions in “Apoptosis” and “Immunity” function types. Interestingly, these two categories, unlike others (such as “Enzymes” or “Transporter”) are mainly associated with protein-protein interactions^95,96^. Hard pathogenic variants that are located at the surface of the protein might have an impact on protein-protein interactions and on the protein function itself. Nonetheless, almost all VEPs tested here do not have any features that could manage these characteristics. Another potential reason is that the VEPs may not have been trained with a sufficient number of hard variants in their training datasets. It ends in models that have limited generalization capabilities for accurately predicting the effects of hard variants.

One important aspect of variant effect prediction worth discussing is the annotation of the variant. Depending on the source of information, the annotation can be of relative quality as we saw with ClinVar. Each database of variant annotation may have their own annotation criterias, which may differ from common ones as it is the case for HGMD databases in which pathogenicity annotations are mainly literature based^6^. Hence, 1000Genomes^2^ variants have also been annotated by crossing database annotations of variants if available. These annotations are accessible through the EnsEMBL genome browser^97^. Users should be aware of how variants are annotated in databases they use, as some annotations might appear human-based but are actually aided by VEPs, potentially introducing bias. While the median AUROC value for ClinVar was 0.93, the same evaluation for 1000Genomes gives only 0.73 (see Figure S10). Although the top-performing methods change a bit, there is a significant reduction in AUROC values.

Finally, we would like to make some suggestions concerning the usability of VEP. One of the main targets of such tools are clinicians who are confronting patients with genetic disease and variants to be analyzed. However, these users are generally far from being experts in the use of bioinformatics tools. Therefore, any VEP should include an “user-friendly” side to be used even by non-specialists. Furthermore, this should be one of the main aims of future developments of VEPs. There are several ways to make the tool easier to use, like developing a web server allowing direct use. Inputs should also be general enough to allow large usability. For example, either genomic coordinates or protein position (or both if possible) could be provided to describe the variant. If the VEP computational time is too important or the VEP is difficult to use, it can be interesting to make available precomputed predictions for all missense mutations possible to avoid the usage of VEPs for non-specialists. This would significantly increase the usability of any VEP regardless of its usage difficulty, such as it is already the case for some of those tested here (such as AlphaMissense, VARITY, VESPA or DeepSav). Moreover, the availability of precomputed predictions is useful for researchers doing benchmarking analysis, like us in this study, as it significantly facilitates the scores retrieving when having various datasets.

Initiatives like the dbNSFP database is a major resource which allows the large diffusion of existing VEP and allows the use of most VEPs used here such as MetaRNN, BayesDel or ClinPred. In addition, it permits their use and the comparison of their predictions totally freely and publicly, which is recommended for an user-friendly use of a global audience.

## Conclusion

In this study, we have made a thorough benchmark analysis on a large set of 65 distinct VEP scores. Amongst them, AlphaMissense performs consistently well on all datasets and is very easy to use as authors have provided precomputed predictions for all possible single amino acid changes in the human proteome. It is actually probably the best option to predict the impact of variants among tested VEPs in this study. However, it is better to use in combination with AlphaMissense, other high performance VEPs (among VARITY, gMVP, MutScore, DeepSAV, CPT, VESPA, SIGMA, EVE, REVEL or MetaRNN) in order to obtain a representative and more nuanced predicted profile of the impact of clinical variants.

We have shown that variants can be categorized into main classes according to their prediction difficulty. easy variants fit quite well with common representation of a variant, meaning that benign variants are at the surface of the protein and pathogenic variants in the protein core. On the other hand, hard variants are characterized in drastically different ways, which may explain their difficulty. Indeed, hard pathogenic variants are located at the surface of the protein whereas hard benign variants are found in the core. This suggests that these pathogenic variants might be hard to predict as they could potentially impact protein-protein interactions; an aspect not efficiently considered by current VEPs.

For future development of VEPs, we recommend aiming for easy usage for non specialist users such as clinicians who are the primary target audience but may not be experts in bioinformatics tools. This can be achieved by developing an user-friendly web-server and/or releasing precomputed predictions for all possible single amino acids substitutions in the human proteome to avoid running the VEP which can be very useful when the actual VEP is quite complicated to use.

## Abbreviations

ACMG: American College of Medical Genetics
AMP: Association for Molecular Pathology
SNP: Single Nucleic acid Polymorphism
OMIM: Online Mendelian Inheritance in Man
HGMD: Human Gene Mutation Database
VEPs: Variant Effect Predictors
AF2: AlphaFold2
ML: Machine Learning
AI: Artificial Intelligence
DMS: Deep Mutation Scanning
T1C: Type 1 Circularity
T2C: Type 2 Circularity
AUROC: Area Under the ROC curve
MCC: Matthews Correlation Coefficient
BER: Balanced Error Rate

## Acknowledgments

The authors would like to thank Mr. Gabriel Cretin, Mr. Yann Vander Meersche and Pr. Catherine Etchebest (DSIMB, Paris, France) for fruitful discussions.

## Author contributions

A.G.dB. and J-C.G have designed and conceptualized the study. R.R. has made the curation of data and all the calculations. Analysis has been made mainly by R.R. all the results have been discussed and improvements have been proposed by all authors. R.R. draws the initial draft and A.G. dB., J-C. G. and J.D. have edited the article. All authors have acknowledged this final version of the manuscript.

## Data and code availability

All datasets and code to obtain predictions are available on GitHub: https://github.com/DSIMB/VEP-Benchmark. The provided pipeline allows users to retrieve 65 prediction scores from standard input formats for variant data, including genomic and protein positions.

## Web Resources

dbNSFPv4.4 (access date: 2023/05/06), http://database.liulab.science/dbNSFP

ClinVar (access date: 2023/12/28), https://www.ncbi.nlm.nih.gov/clinvar/

UniProt (access date: 2023/12/29), https://www.uniprot.org/

Humsavar (access date: 2023/11/08), https://www.uniprot.org/docs/humsavar

SuSPect (access date: 2024/01/07), http://www.sbg.bio.ic.ac.uk/~suspect/

ESM github (access date: 2024/01/04), https://github.com/facebookresearch/esm

AlphaFoldDB (access date: 2024/02/01), https://alphafold.ebi.ac.uk/

Protein Data Bank (access date: 2024/02/01), https://www.rcsb.org/

## Supplementary materials

**Figure S1.**
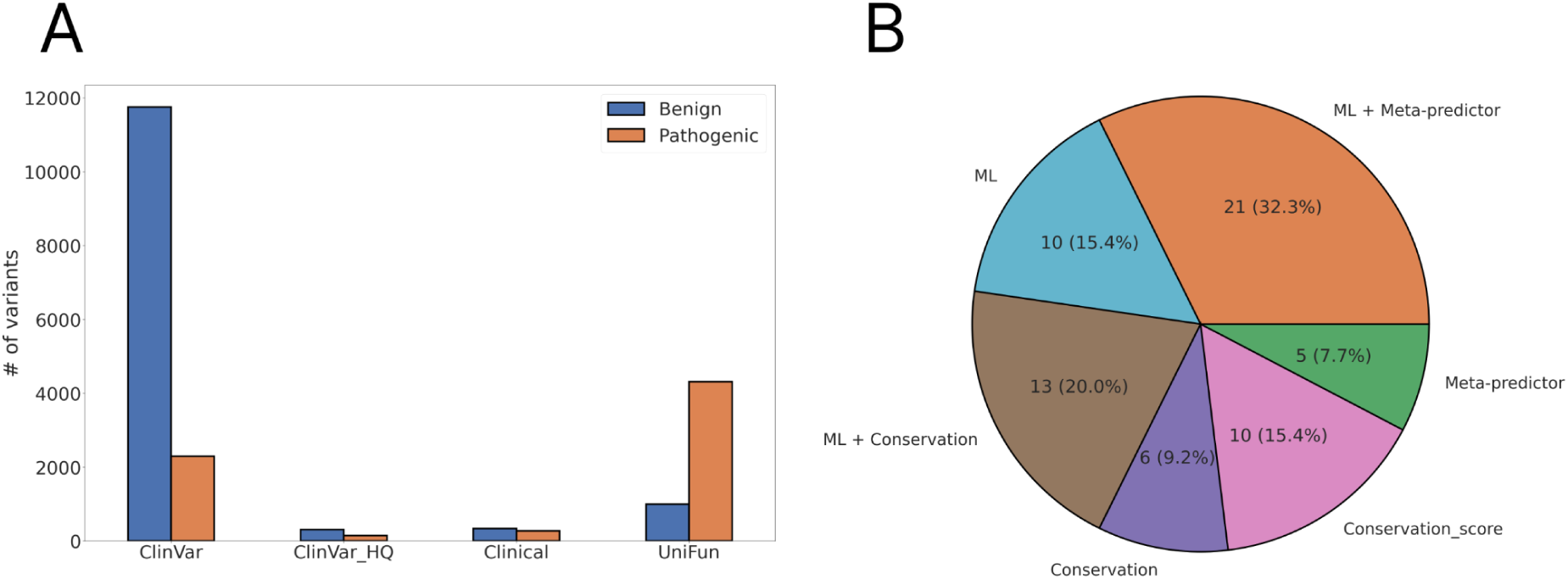
Summary of datasets and VEPs type used. (A) Distribution of benign and pathogenic variants across the different datasets. (B) Amount of VEPs within each VEP group.

**Figure S2.**
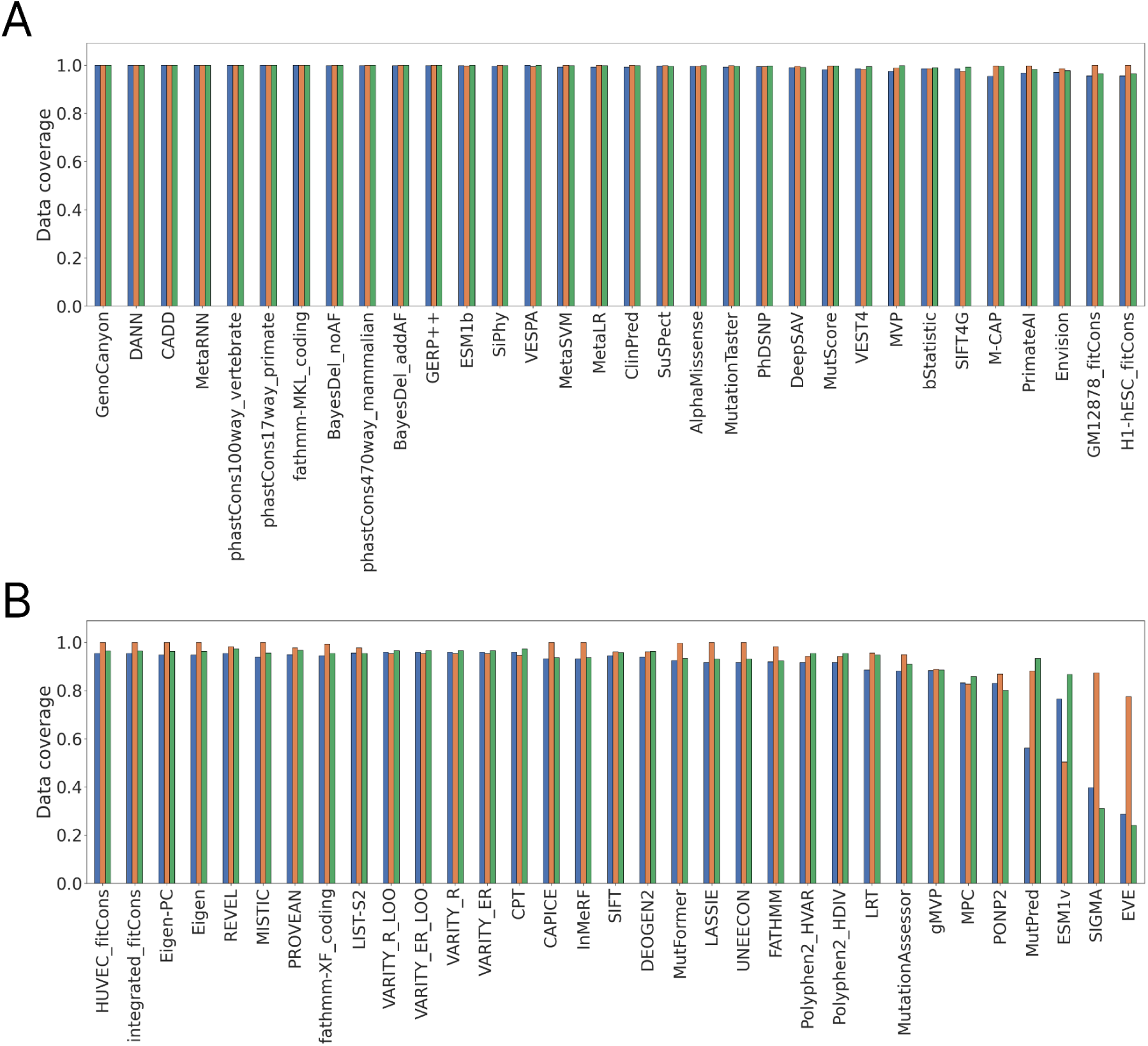
Prediction results availability for each VEP. (A) Data coverage for the first 32 VEPs and (B) for the remaining 35 VEPs are represented for each variant dataset. In blue color ClinVar, in orange clinical dataset and in green UniFun.

**Figure S3.**
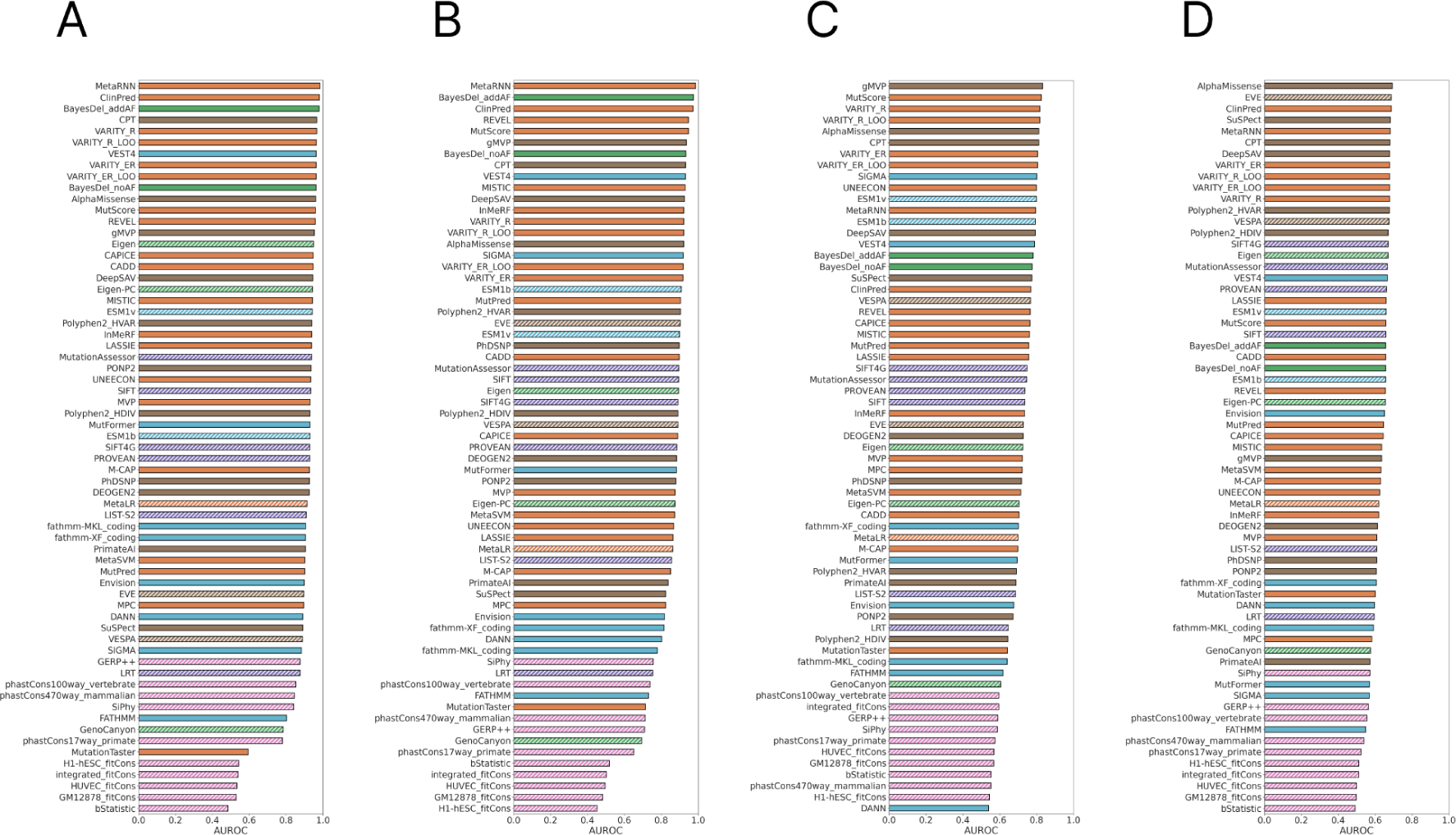
AUROC performances. AUROC values for ClinVar (A), ClinVar_HQ (B), Clinical (C) and UniFun (D). Each bar is colored according to the type of algorithm used by the VEP.

**Figure S4.**
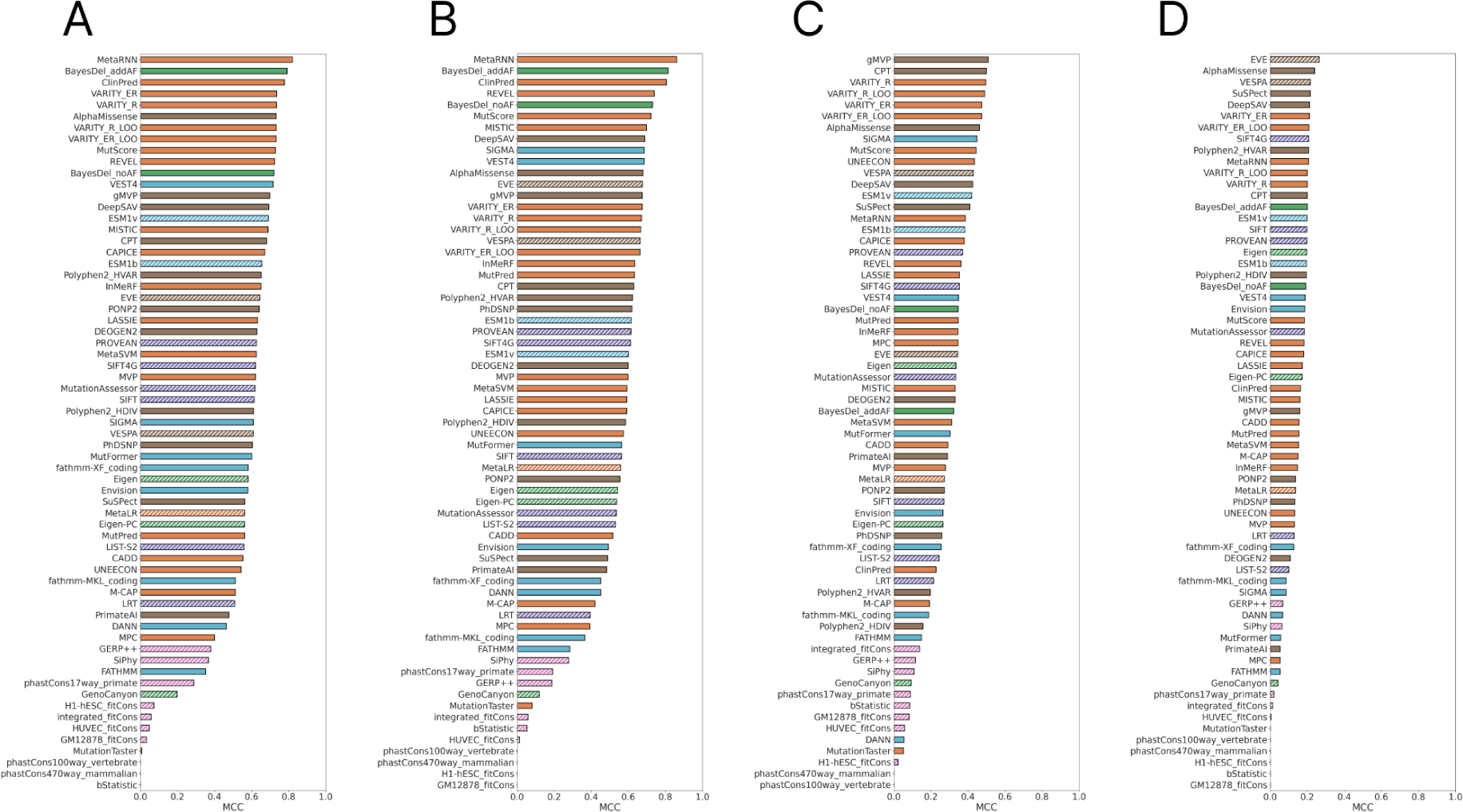
MCC performances. MCC values for ClinVar (A), ClinVar_HQ (B), Clinical (C) and UniFun (D). Each bar is colored according to the type of algorithm used by the VEP.

**Figure S5.**
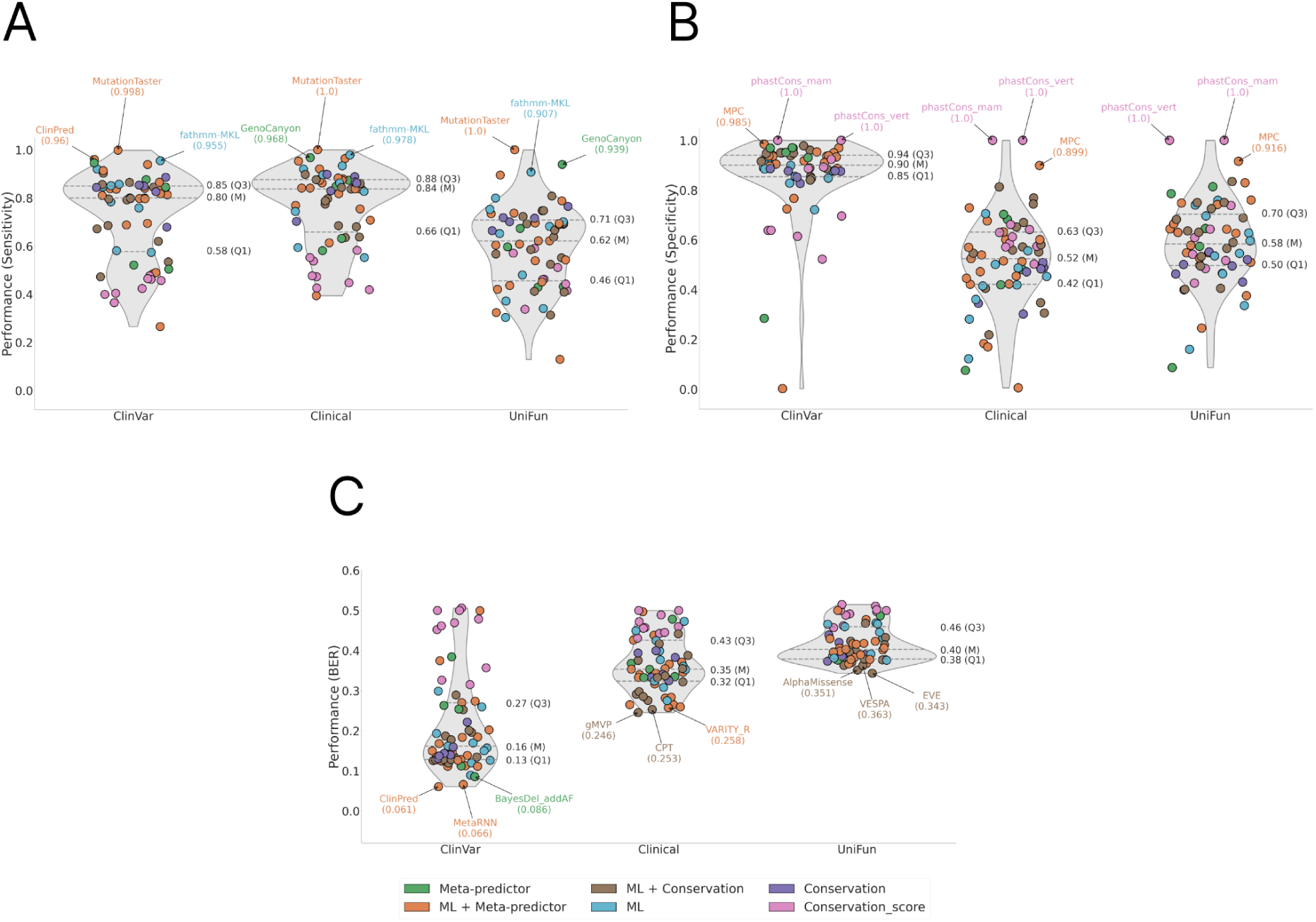
Overview of performance metrics on all datasets. (A) Sensitivity values are represented for each dataset, which measure the accuracy on positive labels. (B) Specificity values are represented for each dataset, which measure the accuracy on positive and negative. (C) BER values are represented for each dataset, which indicate the global error rate in both sensitivity and specificity metrics.

**Figure S6.**
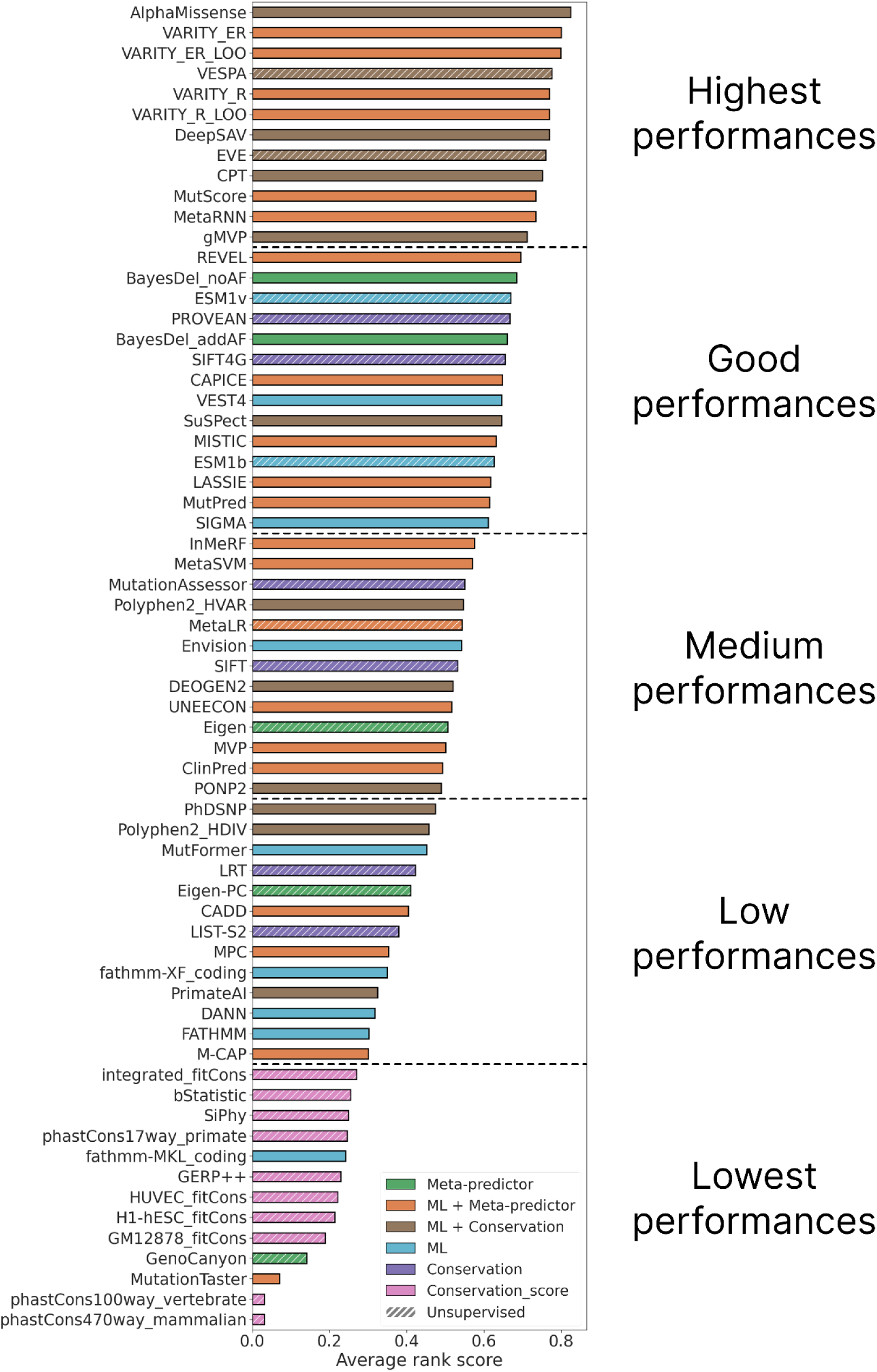
Average rank score and group of performance. The average rank score represents the mean between rank scores values from ClinVar_HQ, Clinical and UniFun datasets and constitute the global performance of each VEP in this study. Several cutoff values are used to regroup VEPs according to their performance. These cutoff values are 0.7, 0.6, 0.4 and 0.3.

**Figure S7.**
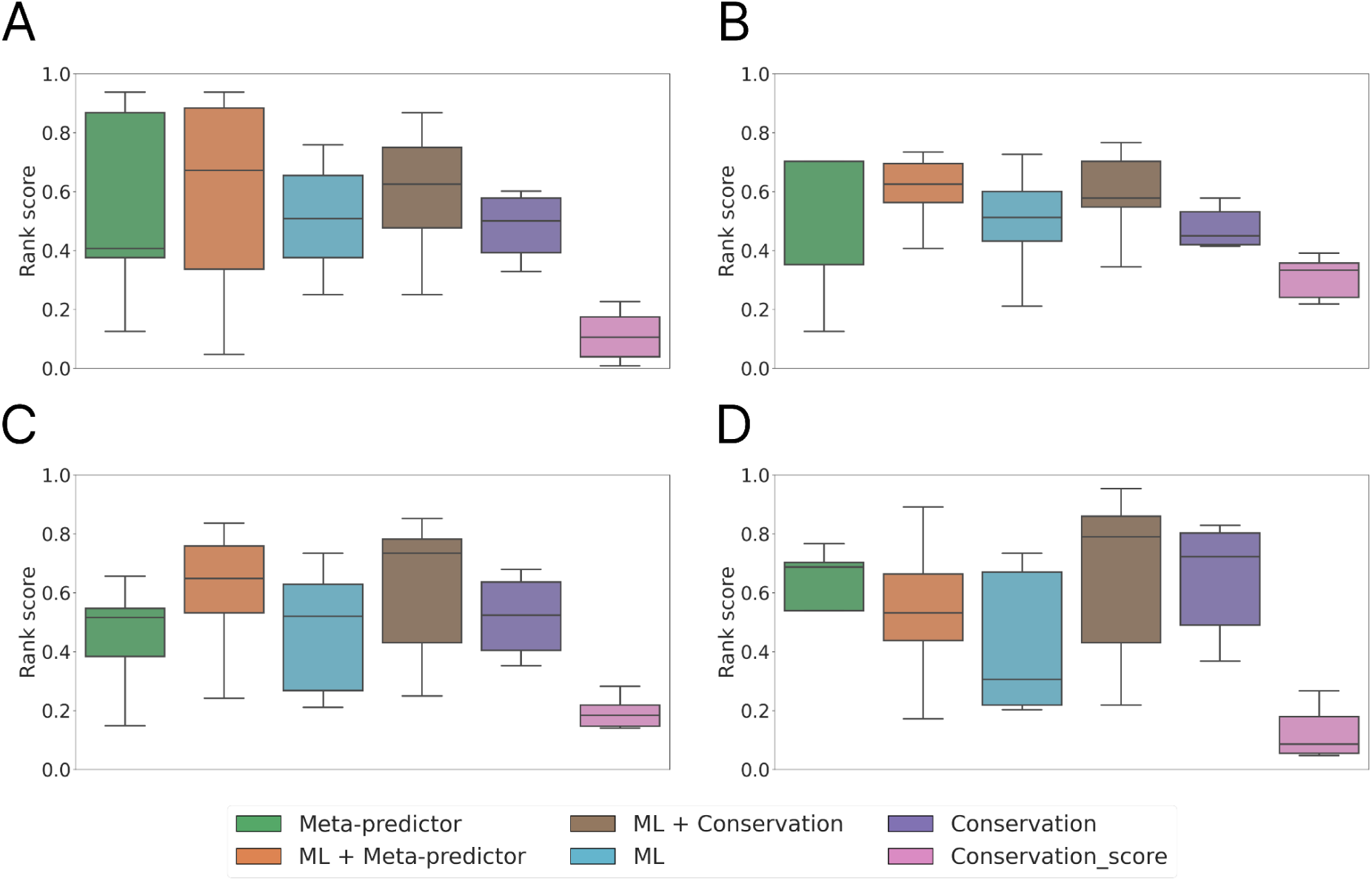
Performance by VEP type. Distribution of rank scores by VEP type on ClinVar (A), ClinVar_HQ (B), Clinical (C) and UniFun dataset (D).

**Figure S8.**
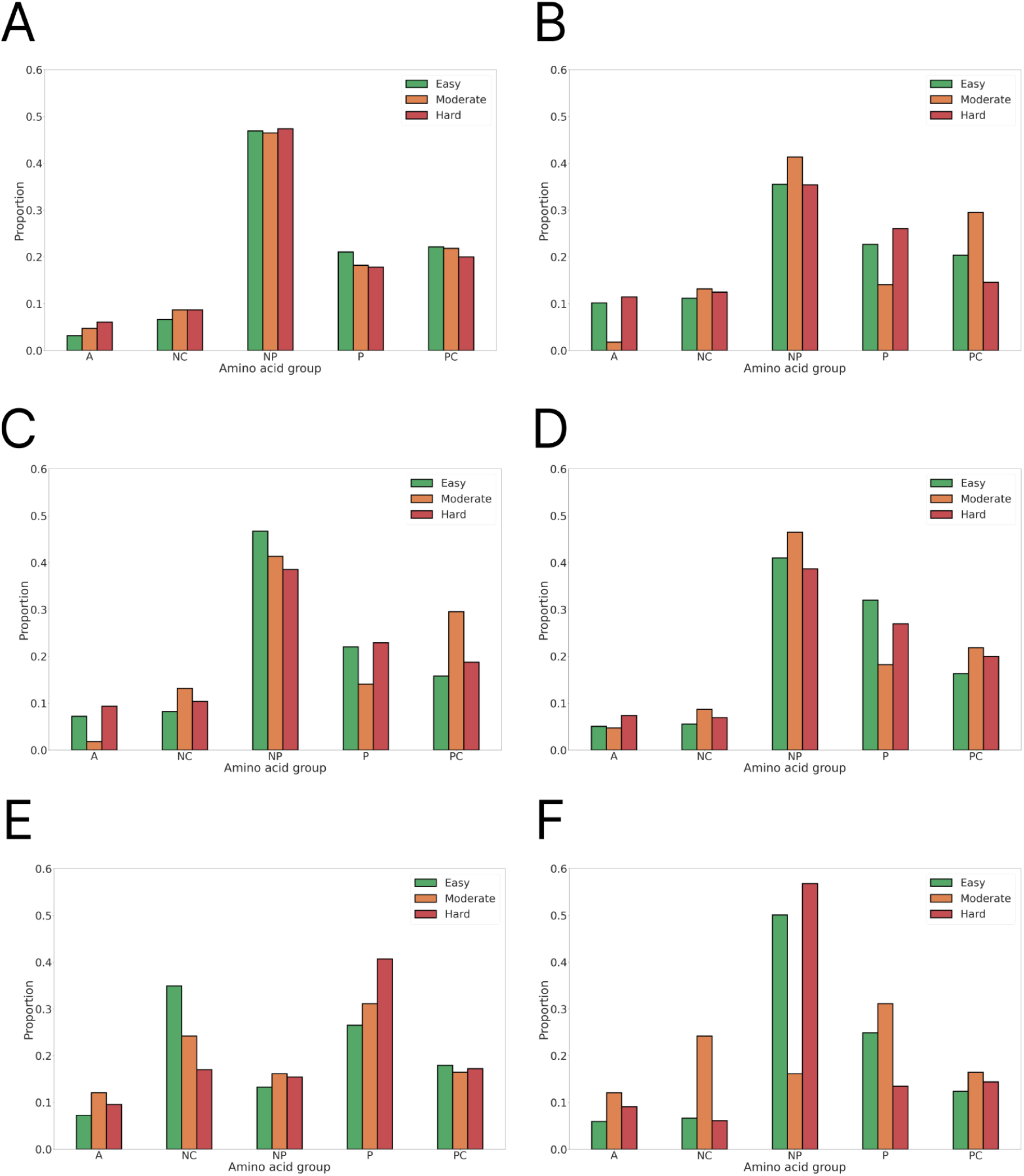
Amino acid composition of easy, moderate, and hard variants. Proportions of wild-type residues in ClinVar (A), Clinical (C), and UniFun (E) datasets, and mutant residues in ClinVar (B), Clinical (D), and UniFun (F) datasets. Amino acids are categorized by physicochemical groups: aromatic (“A”), negatively charged (“NC”), nonpolar (“NP”), polar (“P”), and positively charged (“PC”).

**Figure S9.**
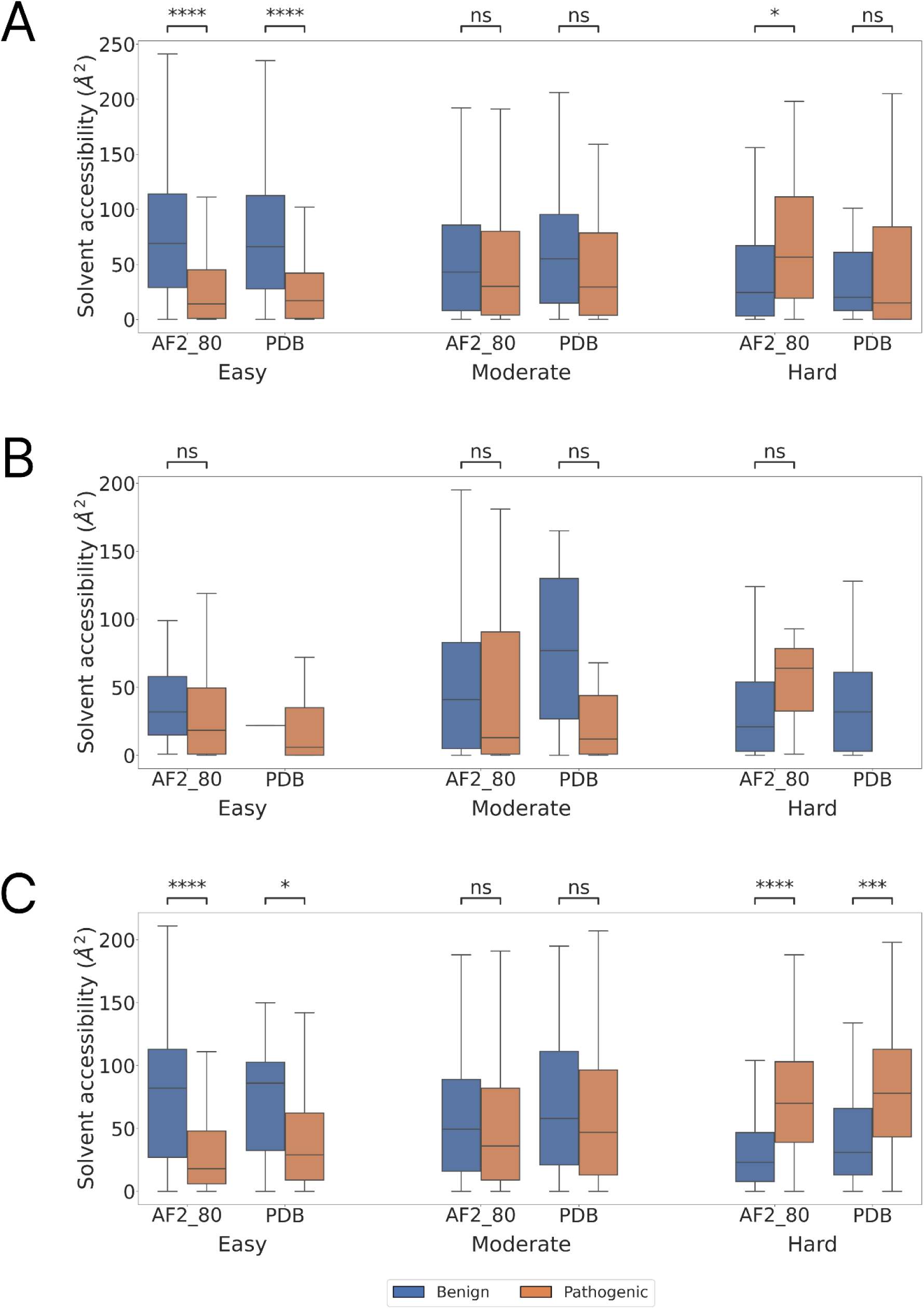
Solvent accessibility analysis. Solvent accessibility values for easy, moderate and hard variants in ClinVar (A), Clinical (B) and UniFun dataset (C). Each panel contains solvent accessibility values for benign (blue) and pathogenic (orange) variants using protein structure from the AF2 database filtered at 80 pLDDT (AF2_80, see methods) or from the PDB (PDB). Mann-Whitney statistical test has been applied for each comparison.

**Figure S10.**
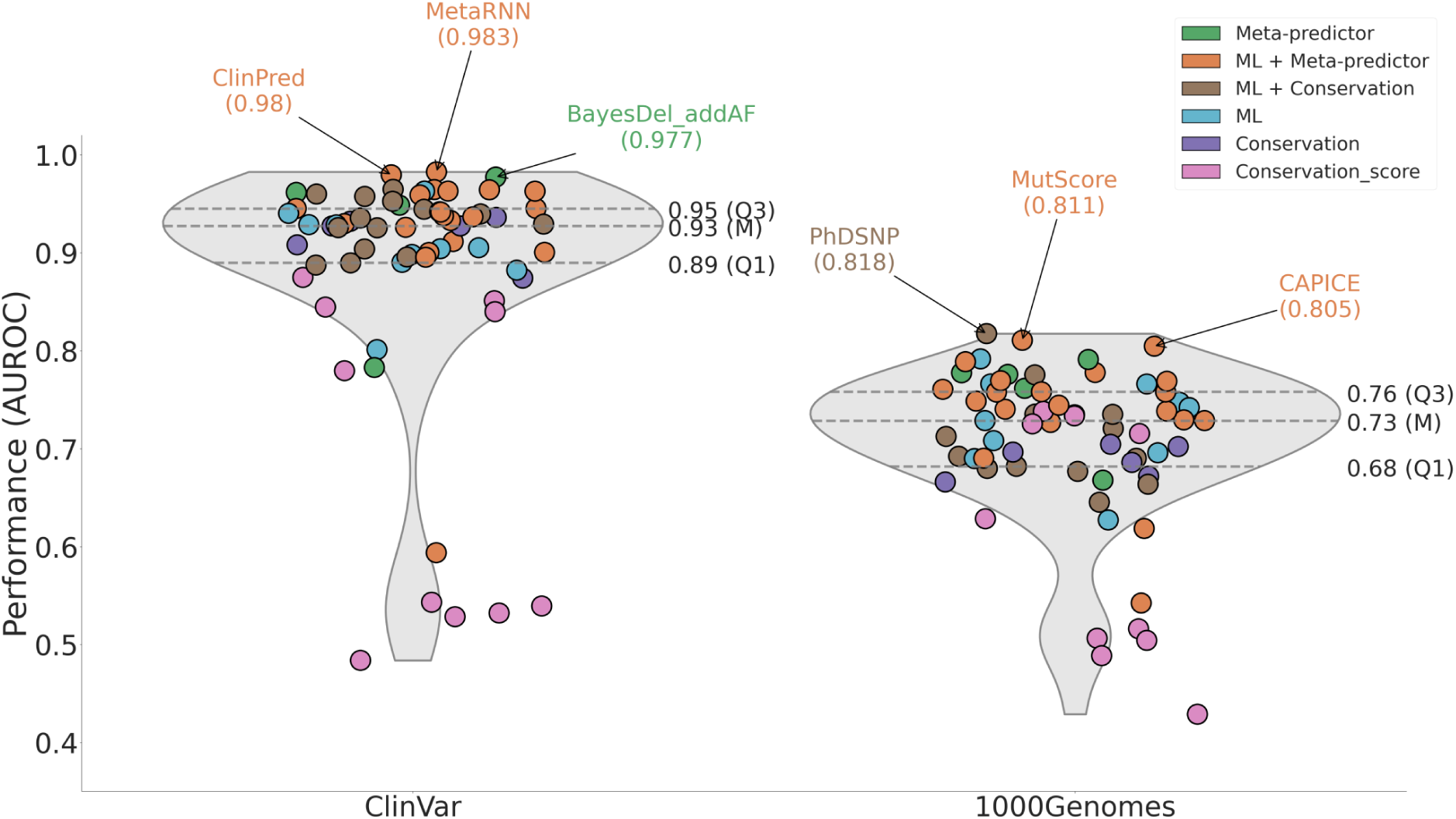
Impact of the dataset. Distribution of AUROC for ClinVar (A) as done in Figure 2A, and 1000Genomes (B).

**Table S1. p-values from pairwise chi-square test comparison between UniProt functions present in easy variants from all datasets.** Values in bold are the ones that are significant according to the corrected alpha from Bonferroni correction in multiple testing analysis. The corrected alpha is equal to 8.3*10^-3 (corresponding to 0.05 divided by 6, the number of tests done for each group).

**Table S2. p-values from pairwise chi-square test comparison between UniProt functions present in moderate variants from all datasets.** Values in bold are the ones that are significant according to the corrected alpha from Bonferroni correction in multiple testing analysis. The corrected alpha is equal to 8.3*10^-3 (corresponding to 0.05 divided by 6, the number of tests done for each group).

**Table S3. p-values from pairwise chi-square test comparison between UniProt functions present in hard variants from all datasets.** Values in bold are the ones that are significant according to the corrected alpha from Bonferroni correction in multiple testing analysis. The corrected alpha is equal to 8.3*10^-3 (corresponding to 0.05 divided by 6, the number of tests done for each group).

**Table S4. Amount of variants for each function group.** Overview of the number of benign and pathogenic variants within each function group across three levels of prediction difficulty: Easy, Moderate, and Hard. Function groups include Apoptosis, Immunity, Metal-binding, Enzyme, Transport, Signal, and Receptor.

**Table S5. Proportion of variants found in each protein function according to variant difficulty.** This table represents the proportions of variants in each function group classified by the difficulty of prediction: Easy, Moderate, and Hard.

